# TBK1 regulates autophagic clearance of soluble mutant huntingtin and inhibits aggregation/toxicity in different models of Huntington’s disease

**DOI:** 10.1101/869586

**Authors:** Ramanath Narayana Hegde, Anass Chiki, Lara Petricca, Paola Martufi, Nicolas Arbez, Laurent Mouchiroud, Johan Auwerx, Christian Landles, Gillian P. Bates, Malvindar K. Singh-Bains, Maurice A Curtis, Richard L. M. Faull, Christopher A. Ross, Andrea Caricasole, Hilal A Lashuel

## Abstract

Phosphorylation of the N-terminal domain of the Huntingtin (HTT) protein (at T3, S13, and S16) has emerged as a key regulator of HTT stability, clearance, localization, aggregation and toxicity. Herein, we report the discovery and validation of a kinase, TANK-binding kinase 1 (TBK1), that specifically and efficiently phosphorylates both wild-type and mutant full-length or N-terminal fragments of HTT *in vitro* (S13/S16) and in cell/ neuronal cultures (S13). We show that overexpression of TBK1 in mammalian cells, primary neurons and a *Caenorhabditis elegans* model of Huntington’s Disease (HD) increases mutant HTTex1 phosphorylation, lowers its levels, increases its nuclear localization and significantly reduces its aggregation and cytotoxicity. Our mechanistic studies demonstrate that the TBK1-mediated neuroprotective effects are due to phosphorylation-dependent inhibition of mutant HTTex1 aggregation and an increase in autophagic flux. These findings suggest that upregulation and/or activation of TBK1 represents a viable strategy for the treatment of HD.

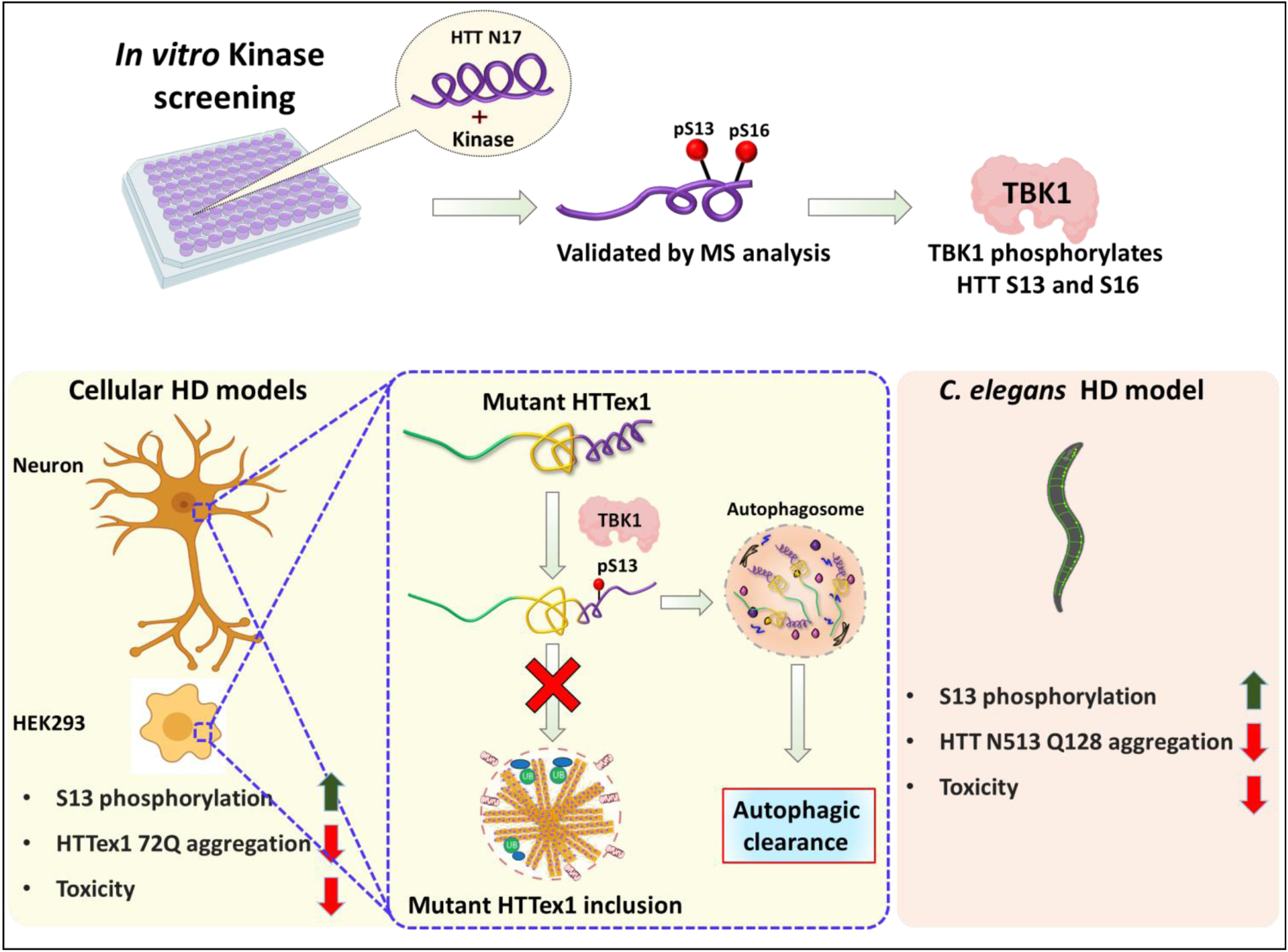
Graphical abstract.

## Introduction

Huntington’s disease (HD) is an autosomal dominant, progressive neurodegenerative disease caused by a CAG triplet repeat expansion (>35) in the Huntingtin gene. This translates into a polyglutamine (polyQ) repeat immediately following the first 17 amino acids (N17) in the Huntingtin protein (HTT) (MacDonald et al., 2003; Myers et al., 1993; Rubinsztein et al., 1996). PolyQ repeats in HTT in the disease range (> 36, hereafter denoted as mutant HTT) render the protein more susceptible to misfolding, thus leading to the formation of HTT aggregates in cells and neurons (Caron et al., 2013; Cui et al., 2014; Daldin et al., 2017; Fodale et al., 2014; Scherzinger et al., 1999). Increasing evidence from human studies has shown that expansion of the polyQ repeat is the causative mutation of HD but not the only contributor to disease onset, duration and severity (Andresen et al., 2007; Lee et al., 2019; Wexler et al., 2004). This suggests that other factors may play important roles in modifying the course of the disease and potentially the patient response to therapies.

Post-translational modifications (PTMs), particularly within the N17 of the HTT protein, has emerged as key regulators of HTT stability, clearance, localization, proteolysis, aggregation and toxicity in different models of HD (Ehrnhoefer et al., 2011; Saudou and Humbert, 2016), including mouse HD models (Gu et al., 2009). Largely these studies involved mutation of PTM-targeted residues to mimic or abolish the presence of relevant PTMs. Among all the putative PTMs that have been identified in the HTT protein, phosphorylation at the N-terminal serine residues S13 and S16 are the most studied for the following reasons: 1) it occurs in close proximity to the polyQ domain and within the N17 domain, which has been shown to play important roles in regulating HTT structure, aggregation and subcellular localization (Gu et al., 2009; Rockabrand et al., 2007); 2) mimicking phosphorylation at these residues in the context of the full-length mutant HTT protein (S13D/S16D) in mice was sufficient to ameliorate HD phenotypes, including motor and psychiatric-like behavioral deficits, mutant HTT aggregation and selective neurodegeneration (Gu et al., 2009); and 3) the levels of phosphorylation at S13/S16 are reduced in HD models (Atwal et al., 2011; Cristina Cariulo, 2019). These observations suggest that targeting HTT PTMs at N17 represents a viable strategy for developing novel disease-modifying therapies for HD. The first step towards assessing the therapeutic potential of targeting N17 PTMs for the treatment of HD is the identification of the enzymes that regulate these PTMs. This is important because increasing evidence shows that phosphorylation-mimicking mutations (e.g., Serine (S) or Threonine (T) to Aspartate (D) or Glutamate (E)) used to investigate the effects of protein phosphorylation did not fully phenocopy the effects of the *bona fide* modifications (Deguire et al., 2018; Paleologou et al., 2008; Szczepanowska et al., 1998) and did not faithfully reproduce the dynamic nature of phosphorylation.

We recently showed that phosphorylation of HTT at S13 and/or S16 inhibits mutant HTTex1 aggregation *in vitro* (Deguire et al., 2018) and increases its conformational flexibility (Daldin et al., 2017). A direct comparison of the effect of phosphomimetics and *bona fide* phosphorylation at these sites also showed that the phosphomimetics (S13D/S16D) only partially reproduced the effect of phosphorylation on aggregation and did not reproduce the effect of single and double phosphorylation at these residues on the helical conformation of N17 (Deguire et al., 2018). Moreover, a recent study using a *Drosophila* model of HD expressing HTTex1 97Q *via* phosphomimetics showed that both S13D and S16D increase aggregation (Branco-Santos et al., 2017), in contrast to observations in mammalian cell models of HD showing that expression of the N-terminal fragments of HTT 1-171 142Q, HTTex1 97Q or HTTex1 82Q with S13D and S16D decreases mutant HTT-induced toxicity and aggregation (Arbez et al., 2017; Atwal et al., 2011; Branco-Santos et al., 2017). Although these findings show that PTMs, such as single or double phosphorylation, are sufficient to modify mutant HTT levels, aggregation, and toxicity, they highlight the limitations of using phosphomimetics. These findings also emphasize the importance of identifying the natural kinases involved in regulating HTT phosphorylation to assess the relevance and therapeutic potential of *bona fide* phosphorylation at these residues.

In the current study, we sought to identify the natural kinases that efficiently phosphorylate HTT at S13 and S16, with the aim of using these kinases to assess the therapeutic potential of phosphorylation at these residues by investigating its effect on mutant HTT aggregation, clearance, and toxicity in cellular and animal models of HD. Using an *in vitro* screen of ∼300 serine and threonine kinases, we identified TANK-binding kinase 1 (TBK1) as one of the most promising candidate kinases. In an *in vitro* phosphorylation assays, TBK1 induced site-specific and quantitative phosphorylation of both wild-type and mutant HTT at both residues S13 and S16. TBK1 is known to regulate the innate immune response and belongs to the IκB kinase family that includes other kinases such as IKKβ, which was previously shown to phosphorylate HTT at these residues (Thompson et al., 2009) as well as at residue T3 (Bustamante et al., 2015), and was recently shown to modulate HTT S13 phosphorylation in HD mice (Ochaba et al., 2019). In this work, the specificity and efficiency of TBK1 for phosphorylating full-length and N-terminal fragments of HTT (HTTex1, HTTN548, HTT-fl) were validated in mammalian cells, primary neuronal culture and *in vivo via* overexpression of the wild-type and kinase-dead variant of TBK1, through a combination of techniques including a novel ultrasensitive immunoassay for pS13 HTT detection (Cristina Cariulo, 2019). Next, we assessed the effect of TBK1-mediated phosphorylation on HTT levels, subcellular localization, aggregation, and toxicity in cellular, neuronal and *Caenorhabditis elegans (C. elegans)* models of HD. In all these model systems, we observed that TBK1 phosphorylated the HTT protein at S13, decreased its levels, inhibited its aggregation and suppressed its toxicity. Furthermore, mechanistic studies showed that these effects were mediated by both TBK1-dependent phosphorylation of HTT, which stabilized the protein and blocked its aggregation, and TBK1-mediated increased autophagic flux, which also promoted the clearance of HTT, leading to a decrease in HTT aggregation. Together, our results show that increasing phosphorylation at S13 could be an effective strategy to reduce mutant HTT aggregation and/or TBK1 activation or upregulation may be a viable therapeutic strategy for the treatment of HD.

## Results

### Identification of TBK1 as a kinase that efficiently phosphorylates HTT at S13 in cellular models of HD

To discover kinases responsible for phosphorylating HTT at S13 and S16, we used a commercial *in vitro* kinase screening assay, which includes ∼300 kinases (Binukumar et al., 2016), using two HTT substrates; a peptide that comprises the first 17 N-terminal residues of HTT (N17), and the first exon of the HTT protein (HTTex1) (Fig. 1A-C). After an extensive *in vitro* validation using the top kinase hits from the screen, which included monitoring of the extent of phosphorylation using mass spectrometry as well as by the use of previously validated phospho-antibodies against T3 (pT3), S13 (pS13) and S16 (pS16) (Bustamante et al., 2015; Cristina Cariulo, 2019; Deguire et al., 2018), TBK1 was identified as the only kinase that selectively and robustly phosphorylated HTTex1 *in vitro* at S13 and S16 (Fig. 1C). Additionally, TBK1 mediated the efficient and site-specific phosphorylation of recombinant longer N-terminal HTT fragments at S13 and S16 *in vitro* (Supplemental Fig. 1A-B). We also observed that phosphorylation by TBK1 was significantly faster than the previously identified S13/S16 HTT kinase, IKKβ (Bustamante et al., 2015; Thompson et al., 2009) at both sites (Supplemental Fig. 2A, D). Collectively, these findings demonstrate that TBK1 efficiently and site-specifically phosphorylates recombinant HTT at S13 and S16 *in vitro,* suggesting that it could be one of the natural kinases that may regulate HTT phosphorylation and cellular properties *in-vivo*.

**Figure 1:**
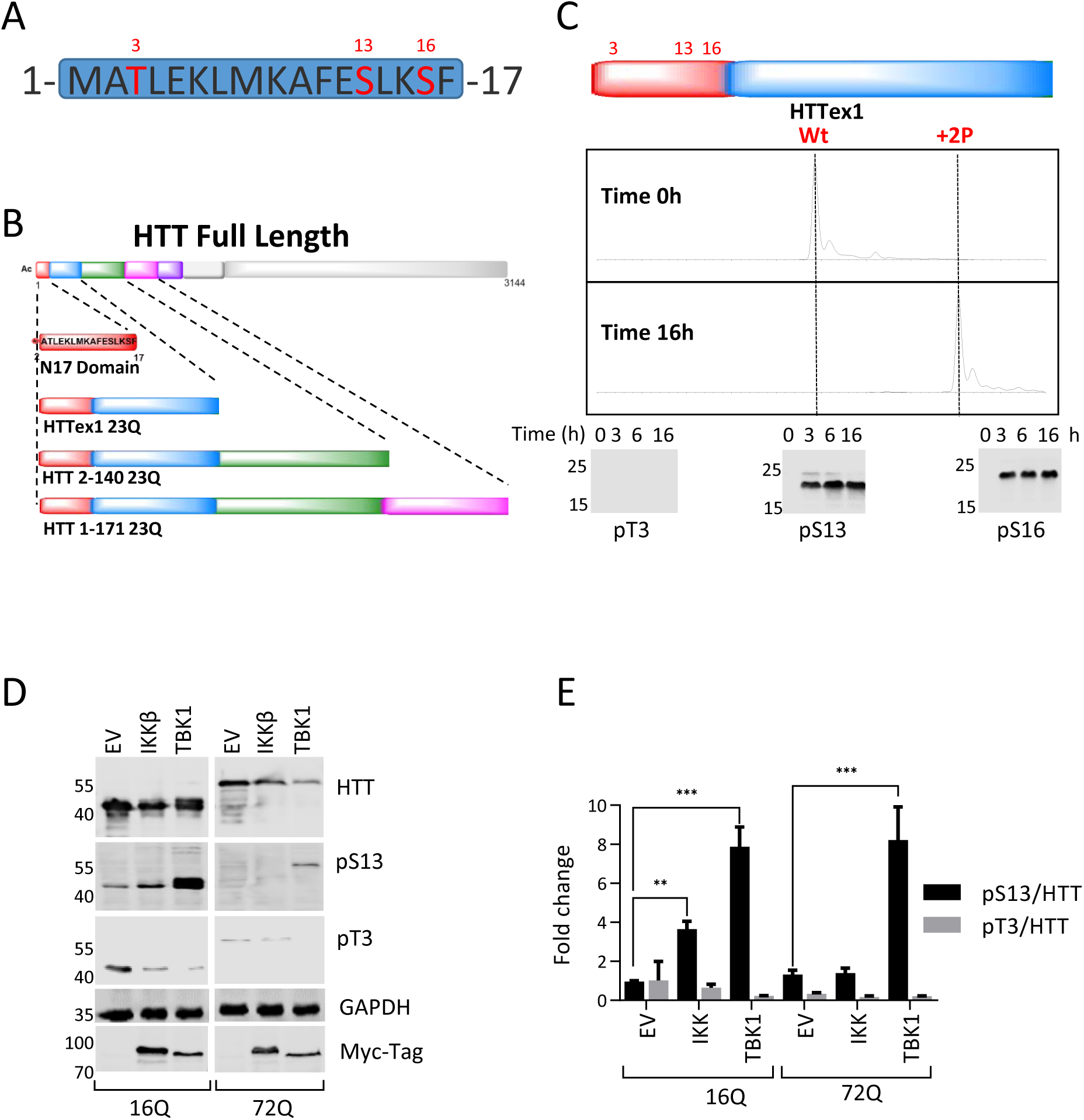
Identification of TBK1 as a kinase that efficiently phosphorylates HTT at S13 in cellular models of HD: (**A**) Potential Phosphorylation sites in the HTT N17 domain (T3, S13 and S16). (**B**) List of all the different substrates used for the *in vitro* kinase validation: Nt17, HTTex1 and HTT longer fragments. (**C**) Mass spectra of recombinant HTTex1 23Q after 16 hrs of co-incubation with by TBK1 showing complete phosphorylation of S13 and S16. The lower panel is a representative western blot of the same phosphorylation reaction (upper panel) using anti-pT3, - pS13 and -pS16 specific antibodies after the indicated phosphorylation reaction times. **(D)** Representative western blot of HTT and HTT pS13 upon co-expression of HTTex1 16Q and 72Q eGFP with the indicated kinases for 48 hrs in HEK293T cells. **(E)** Quantification of the fold changes in the HTT pS13 and pT3 ratios to total HTT compared to empty vector (EV) control upon co-expression of the indicated kinase from the experiments like in D.

Next, to determine whether TBK1 phosphorylates the N-terminal domain of HTT in cells, we assessed the level of phosphorylation at S13 upon co-expression of wild-type (16Q) or mutant (72Q) HTTex1 with TBK1 in HEK293 cells and assessed phosphorylation by western blot (WB) using the phospho-antibodies against T3 (pT3), S13 (pS13). We observed that TBK1 co-expression for 48 hours led to a strong ∼5-10-fold increase in phosphorylation at S13 (Fig. 1D-E) compared to empty vector (EV) control. Before this study, IKKβ was the only kinase that was reported to phosphorylate HTT at S13/S16 with a preference for HTT containing unexpanded polyQ repeats (Thompson et al., 2009). Therefore, we sought to directly compare the phosphorylation efficiency of TBK1 and IKKβ on mutant and wild-type HTTex1. As shown in Figure 1 D and E, IKKβ indeed phosphorylated HTTex1 16Q more efficiently than 72Q, and the phosphorylation levels were much lower than the levels achieved upon co-expression with TBK1, consistent with our *in vitro* data (Supplemental Fig. 2A-D). In our *in vitro* assay, we observed that TBK1 phosphorylated HTT at both S13 and S16, but due to the lack of a specific antibody that recognizes phosphorylation at S16 in cell lysates, we could not directly evaluate whether TBK1 phosphorylated S16 in cellular models by WB. To confirm the sites of phosphorylation in cells, we performed MS/MS analysis of HTTex1 immunoprecipitated from TBK1 and TBK1 KD co-expressing wild-type 16Q HTTex1 HEK293 cells. While we detected the phosphorylation at S13, we did not observe any phosphorylation at S16 on HTT under these conditions (Supplemental Fig. 3 A-B). These results suggest that TBK1 phosphorylates HTT at S13 but not at S16 in cells, whereas both sites *in vitro* (Fig. 1C).

Next, we asked whether the kinase activity of TBK1 was required for the phosphorylation of HTT. To answer this question, we compared the level of phosphorylation at S13 upon co-expression of HTTex1 16Q or 72Q with wild-type TBK1 or a TBK1 kinase-dead (TBK1 KD) mutant (K38A) (Richter et al., 2016). While WT TBK1 increased S13 phosphorylation, no significant increase in phosphorylation was detected upon expression of TBK1 KD (Fig. 2A-B). This confirms that HTT S13 phosphorylation is dependent on the catalytic activity of TBK1. Next, we sought to confirm the site-specificity of TBK1 phosphorylation of HTT in cells. To this end, we co-expressed HTT with alanine mutations at residue 3 (T3A) or residues 13/16 (S13A/S16A) with TBK1 or TBK1 KD. We observed that TBK1 did not phosphorylate HTTex1 S13A/S16A but phosphorylated S13 on HTTex1 T3A (Fig. 2A-B), confirming that TBK1 specifically phosphorylates S13 on HTTex1. We also observed the interaction of TBK1 with both 16Q and 72Q HTTex1 or HTTex1 S13A/S16A when we co-immunoprecipitated HTT from the cell lysates (Supplemental Fig. 3C-D). Interestingly, in contrast to previous reports (Thompson et al., 2009) and our observations for IKKβ in this study, TBK1 phosphorylated both wild type and mutant HTTex1 with the same efficiency (Fig. 1D-E and Fig 2A-B). These findings suggest that the polyQ expansion does not significantly influence TBK1-dependent phosphorylation of S13.

**Figure 2:**
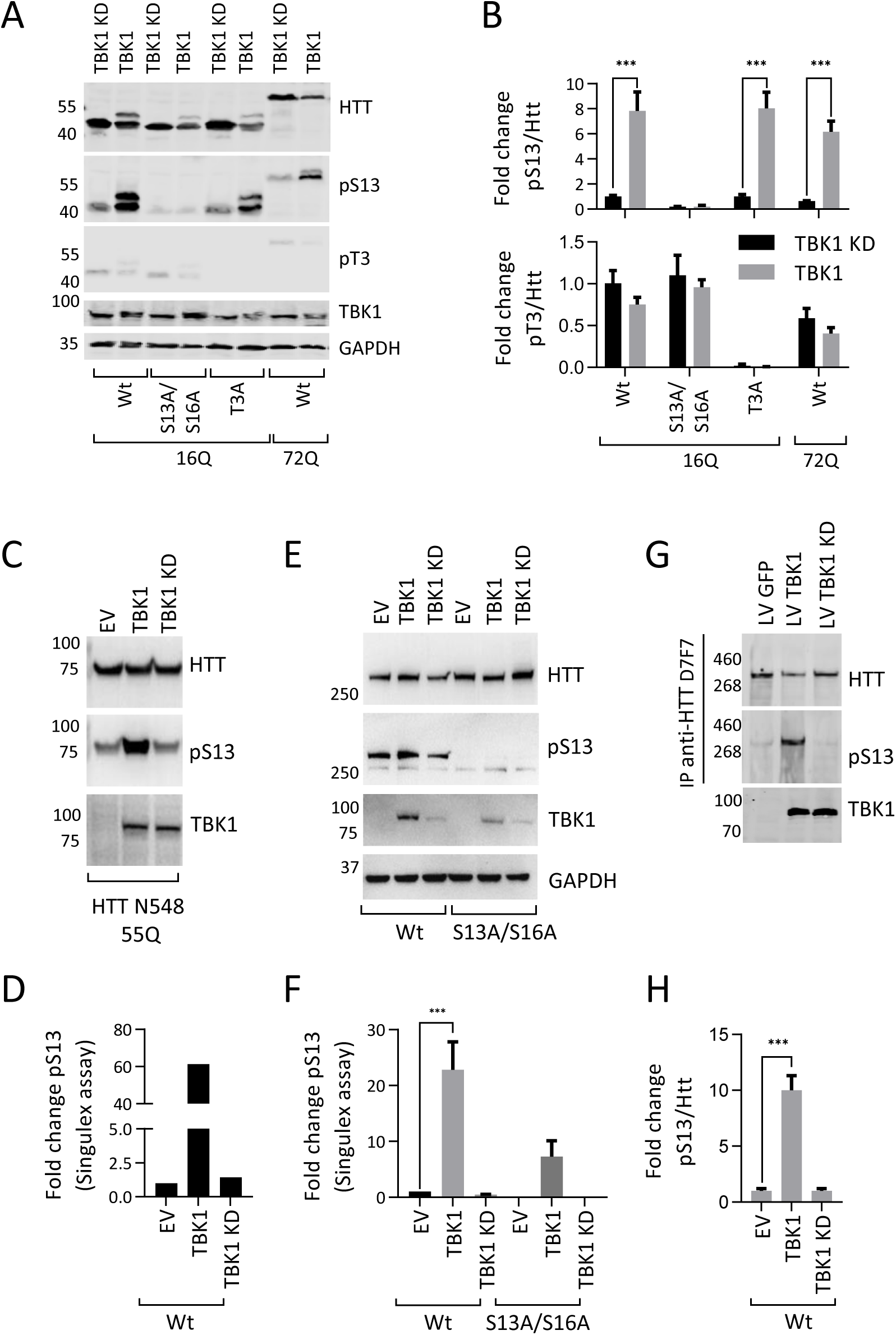
TBK1 phosphorylates HTT in cellular models: (**A**) Representative western blot of HTT, HTT pS13 and HTT pT3 upon co-expression of HTTex1 16Q eGFP, mutants with S to A substitutions at T3 or S13/S16 or HTTex1 72Q eGFP with TBK1 or TBK1 kinase dead (KD) for 48 hrs in HEK293T cells (the additional high molecular weight HTTex1 and HTT pS13 band is possibly due to the co-occurrence of phosphorylation at both S13 and T3) **(B)** Quantification of the fold changes in the HTT pS13 (upper panel) and pT3 (upper panel) ratio to total HTT compared to those of the kinase dead mutant from the experiments in A. **(C)** Representative western blot of HTT and HTT pS13 upon co-expression of HTT N543 16Q with the indicated kinase for 48 hrs in HEK293T cells. **(D)** Quantification of HTT pS13 using the Singulex assay upon co-expression of HTT N543 16Q with the indicated kinase for 48 hrs in HEK293T cells from the experiments in C. (**E**) Representative western blot of HTT and HTT pS13 upon co-expression of HTT FL 23Q or 48Q with the indicated kinase for 48 hrs in HEK293T cells. **(F)** Quantification of HTT pS13 using the Singulex assay upon co-expression of HTT FL 23Q or 48Q with the indicated kinase for 48 hrs in HEK293T cells from the experiments in E. **(G)** Representative western blot of immunoprecipitated (IP) HTT, HTT pS13 and HTT pT3 after lentivirus (LV)-mediated overexpression of TBK1 or TBK1 KD for 72 hrs in rat primary striatal neuronal cells. **(H)** Quantification of the fold changes in the HTT pS13 and pT3 ratios to total HTT compared to those of the kinase dead mutant from the experiments like in G.

Under physiological conditions, HTT exists as a mixture of different N-terminal fragments and full-length HTT (Mende-Mueller et al., 2001). Therefore, we sought to examine whether TBK1 could phosphorylate longer N-terminal fragments and full-length HTT proteins. We co-expressed either the mutant HTTN548 (Q55) fragment or a full-length mutant HTT (Q48) with TBK1 or TBK1 KD in HEK293T cells and analyzed S13 phosphorylation by WB using an HTT pS13 antibody and using a new, sensitive Singulex assay, both of which we recently reported (Cristina Cariulo, 2019). We observed by WB that TBK1 phosphorylated both the HTTN548 Q55 fragment (Fig. 2C) and full-length Q48 at S13 (Fig. 2E). Quantitative Singulex immunoassay analysis of cell lysates showed that TBK1 co-expression led to approximately 60- and 20-fold increases in phosphorylation at S13 of HTTN548 (Fig. 2D) and HTT-fl Q48 (Fig. 2F), respectively. As expected, we did not observe any significant increase in phosphorylation upon co-expression of TBK1 or TBK1 KD with HTT-fl 48Q bearing the S13A and S16A mutations (Fig. 2F).

We further assessed whether TBK1 could phosphorylate endogenous HTT in neurons. We overexpressed TBK1 or TBK1 KD using lentivirus-mediated transduction in rat primary striatal neuronal cultures. Striatal medium spiny neurons are most affected in HD (Ehrlich, 2012) and striatal cells have been used as a model for HD studies (Trettel et al., 2000). We immunoprecipitated endogenous HTT from these striatal culture lysates using HTT antibody (D7F7) followed by immunoblotting with an anti-pS13 antibody. We observed an increased level of HTT pS13, but not pT3 upon TBK1 expression. This confirmed that TBK1 phosphorylated endogenous HTT at S13 (Fig. 2G-H). Next, to investigate whether TBK1 is one of the endogenous enzymes involved in regulating the levels of endogenous phosphorylation of HTT at S13, we used a genetic knockdown or pharmacological inhibition of TBK1 in primary neurons or *Tbk1* -/-, *Tnf-α* -/- mouse model (Hemmi et al., 2004; Taniguchi et al., 1997) primary neurons. We could not observe any changes in HTT pS13 levels upon lowering TBK1 levels or inhibiting its activity (Supplemental Fig. 3E-J). Despite this, we still observed an increase in HTT levels under these conditions. These data collectively indicate that TBK1 regulates HTT levels, but may not be the only kinase involved in the regulation of HTT phosphorylation at S13.

### TBK1 overexpression affects HTT subcellular localization and reduces its aggregation and cytotoxicity

It has been reported that the first 17 amino acids of HTT have multiple roles in regulating the cellular properties of HTT, including its subcellular localization (Atwal et al., 2007; Rockabrand et al., 2007; Steffan et al., 2004). HTT N17 domain was shown to contain a translocated promoter region (TPR)-dependent nuclear export signal (Atwal et al., 2007; Benn et al., 2005; Cornett et al., 2005; Maiuri et al., 2013; Rockabrand et al., 2007; Steffan et al., 2004). Mimicking phosphorylation at S13 and S16 (S13D/S16D) increased the localization of the HTT N17 peptide, HTTex1 or longer N-terminal HTT fragments to the nucleus (Arbez et al., 2017; Atwal et al., 2011; Havel et al., 2011; Thompson et al., 2009; Zheng et al., 2013). Moreover, we recently reported that single or double phosphorylation at S13 and/or S16 of HTTex1 significantly enhanced the targeting of mutant HTTex1 fibrils to the nuclear compartment (Deguire et al., 2018). As TBK1 increased S13 phosphorylation, we hypothesized that this phosphorylation could also increase the nuclear localization of HTT. To test this hypothesis, we co-expressed wild-type or mutant HTTex1 and TBK1 or TBK1 KD in HEK293T cells and then comparatively assessed the levels of nuclear and cytosolic HTT by immunocytochemistry (ICC). We observed the increased nuclear intensity of wild-type or mutant HTTex1 eGFP upon TBK1 co-expression (Supplemental Fig. 4A-B). We also assessed the levels of HTT proteins biochemically in purified nuclear and cytosolic fractions. Confirming the imaging results, nuclear levels of wild-type or mutant HTTex1 were increased upon TBK1 co-expression (Supplemental Fig. 4C-D). In summary, TBK1 phosphorylation of HTT increased the nuclear localization of HTT in line with previous observations (Atwal et al., 2011; Thompson et al., 2009).

Previous studies indicated that phosphorylation or the introduction of phosphomimetic mutations at both S13 and S16 (Anne et al., 2007; Arbez et al., 2017; Branco-Santos et al., 2017; Chiki et al., 2017; Deguire et al., 2018) could suppress mutant HTT aggregation *in vitro* and in cellular models. However, the effects of *bona fide* phosphorylation on mutant HTTex1 aggregation were evaluated only *in vitro* using semisynthetic proteins (Daldin et al., 2017; Deguire et al., 2018). Therefore, we tested whether TBK1-induced phosphorylation affected the aggregation of mutant HTT in HEK293 cells. We used HTTex1 72Q eGFP as a surrogate model of mutant HTT aggregation because the overexpression of this protein has consistently been shown to lead to extensive inclusion formation in different mammalian cells (Arrasate et al., 2004). We co-expressed either phosphorylation-deficient T3A, S13/S16A or phosphorylation-competent HTTex1 containing 72Q or 16Q repeats with TBK1 in HEK293T cells for 48 hours. Then, we performed a WB analysis of soluble protein extracts and a filter retardation assay to assess the level of aggregated mutant HTT in the insoluble fraction. In parallel, we also evaluated the number of TBK1-expressing cells presenting cytoplasmic and nuclear aggregates using ICC. Figure 3A and B show that TBK1 co-expression significantly reduced the levels of soluble wild-type or mutant HTTex1 in an HTT S13/S16 phosphorylation-independent manner, as evidenced by the decrease in the levels of HTTex1 in mutants in which phosphorylation at T3 (T3A) or S13/S16 (S13A/S16A) was prevented by mutation to alanine (Fig. 3A-B). Interestingly, we observed a stronger effect for TBK1 than for IKKβ on lowering both soluble wild-type and mutant HTTex1 levels. The filter retardation assay also showed a phosphorylation-independent reduction in HTTex1 72Q aggregates in the insoluble cellular fraction (Fig. 3C-D). Using ICC, we observed that 35-45% of cells expressing phosphorylation-competent HTTex1 72Q or the phosphorylation-deficient mutants S13/S16A and T3A formed aggregates, whereas only ∼10% of cells co-expressing the same constructs with TBK1 showed aggregate formation (Fig. 3E-F). A similar reduction in soluble and aggregated mutant HTT levels was observed for the phosphorylation-deficient HTTex1 mutants S13A/S16A and T3A (Fig. 3E-F). These results indicate that the TBK1-induced reduction of soluble HTTex1 and HTT aggregates could be mediated by mechanisms not entirely dependent on phosphorylation of N17 at S13. HEK293T cells transfected with HTTex1 72Q showed nuclear aggregates in approximately 10% of cells at the indicated time point.

**Figure 3:**
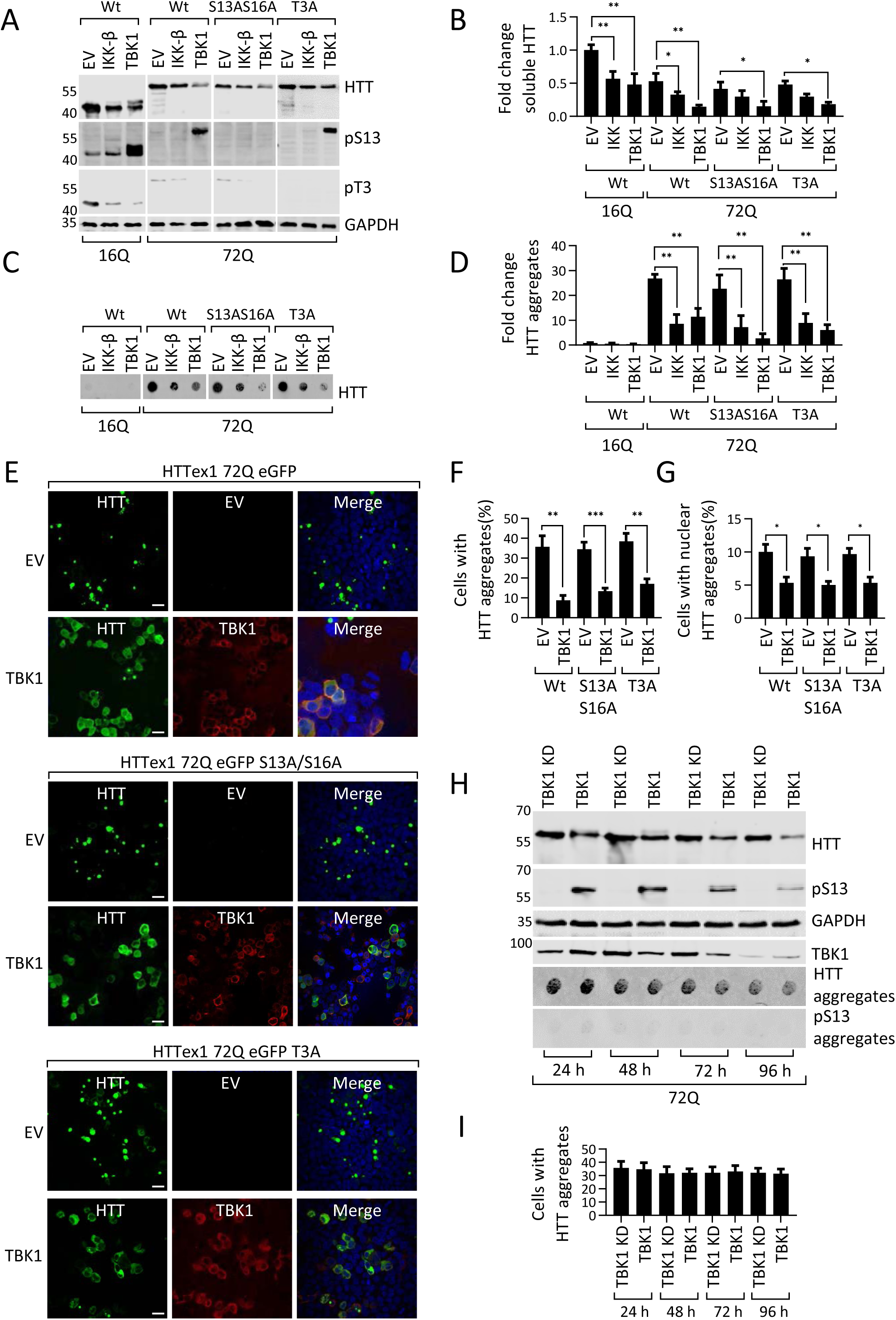
TBK1 overexpression affects subcellular localization and reduces aggregation and cytotoxicity of HTT: **(A)** Representative western blot of soluble HTT, HTT pS13 and HTT pT3 upon co-expression of HTTex1 16Q eGFP or 72Q eGFP or its phosphorylation-incompetent mutants with the indicated kinases for 48 hrs in HEK293T cells. **(B)** The graph indicates the fold change in HTT in samples like illustrated in A, relative to empty vector controls normalized to GAPDH. **(C)** Representative immunoblot of the HTT filter retardation from the insoluble cellular fraction upon expression of HTTex1 16Q eGFP or 72Q eGFP or its phosphorylation-incompetent mutants with the indicated kinases for 48 hrs in HEK293T cells. **(D)** Quantification of the fold change in HTT aggregates compared to EV expressing HTTex1 16Q from the experiments in like in C**. (E)** Representative immunofluorescence images of HTT aggregates upon co-expression of HTTex1 72Q eGFP or its phosphorylation-incompetent mutants with the indicated kinases for 48 hrs in HEK293T cells (scale bar 20μM). **(F)** Quantification of the percentage of co-transfected cells presenting aggregates upon expression of the indicated kinase, from the experiments like in **E**. (**G**) Quantification of the percentage of cells presenting nuclear aggregates from the experiments like in E. **(H)** Representative western blots of the HTTex1 72Q eGFP soluble fraction and aggregates from HEK293T cells transfected first with HTTex1 72Q eGFP and then after 48 hrs with TBK1 to enable formation of aggregates prior to kinase expression; cells were lysed at the indicated time. (**I**) Quantification of the percentage of cells presenting aggregates and kinase expression from the experiments in H.

Therefore, we further examined the effect of TBK1 co-expression on nuclear aggregates. We observed that TBK1 expression also reduced the level of nuclear aggregates formed by HTTex1 72Q eGFP in these cells (Fig. 3G). Interestingly, we also observed that similar to TBK1, IKKβ could decrease HTTex1 levels and mutant HTTex1 aggregates in a phosphorylation-independent manner (Fig. 3C-D).

Next, we sought to determine whether the decrease in the levels of soluble and aggregated mutant HTTex1 was dependent on TBK1 kinase activity. To test this hypothesis, we co-expressed phosphorylation-competent HTTex1 72Q and phosphorylation-deficient S13/S16A or the T3A mutant with TBK1 or its TBK1 KD in HEK293T cells for 48 hours. WB analysis of soluble protein extracts and quantification of aggregated mutant HTT by a filter retardation assay showed that TBK1 but not TBK1 KD decreased soluble and aggregated HTTex1 72Q (Supplemental Fig. 4E-F). In summary, our results establish that TBK1 co-expression lowers HTT levels in an S13/S16 phosphorylation-independent manner, suggesting that TBK1 may be involved in regulating HTT clearance largely *via* other mechanisms.

To determine whether TBK1-induced reduction of HTT aggregate formation could be mediated by its interaction or phosphorylation of HTT aggregates or inclusions, we assessed whether TBK1 or IKKβ could phosphorylate *in vitro-*generated HTTex1 fibrils or mutant HTTex1 aggregates in cells. Towards this aim, we first assessed the ability of TBK1 and IKKβ to phosphorylate preformed HTTex1 43Q fibrils in an *in vitro* assay and monitored the extent of phosphorylation at S13 by filter retardation immunoblot and mass spectrometry analysis. Interestingly, we observed that neither TBK1 nor IKKβ could phosphorylate preformed HTTex1 43Q fibrils, suggesting that S13 residue is readily not accessible in the fibrillar conformation or kinase was not able to interact with HTTex1 fibrils (Supplemental Fig. 4G). We also did not observe any disaggregation or release of HTTex1 43Q monomers during the *in vitro* phosphorylation reaction as evidenced by the absence of change in the levels of HTTex1 43Q fibrils on the filter retardation assay (Supplemental Fig. 4G).

To determine whether TBK1 could phosphorylate already formed HTT aggregates in cells, we expressed mutant HTTex1 72Q in HEK293T cells for 48 hours. At this time point, many cells showed mutant HTTex1 aggregates. Then, we transfected TBK1 and monitored the levels of soluble HTT, HTT aggregates, and pS13 HTT over time by WB and filter retardation assay. First, we did not observe phosphorylation at S13 in the HTTex1 72Q aggregates retained in the filter retardation assay (Fig. 3H). Second, there was no increase in the pS13 levels of HTT aggregates upon overexpression of TBK1 after HTT aggregate formation (Fig. 3H). These observations suggest that TBK1 phosphorylates monomeric or soluble HTT but not fibrils or inclusions of mutant HTTex1 in a cellular context, consistent with the data obtained in our *in vitro* phosphorylation assays (Supplemental Fig. 4G).

Next, we examined whether TBK1 expression promoted the clearance of aggregated mutant HTT. Consistent with our observation in Figure 3A and B, we observed that soluble HTT levels were reduced and S13 phosphorylation was increased upon TBK1 expression at various time points (Fig. 3H). The levels of aggregates were not reduced upon TBK1 overexpression at any given time point, suggesting the limited role of TBK1 in the phosphorylation or clearance of mutant HTT aggregates once they were formed. We observed a decrease in soluble HTT and kinase levels and HTT aggregates at later time points, possibly due to the transient transfection. Accordingly, when testing whether TBK1 could induce clearance of HTT inclusions by ICC, we observed no reduction in the number of HTT inclusions upon overexpression of TBK1 (Fig. 3I). These results indicate that the TBK1-induced reduction in HTT levels and aggregate formation is mediated by TBK1 kinase activity-dependent processes that act at the level of soluble HTT or during the early stages of its oligomerization, before HTT fibrillization and inclusion formation. In line with this hypothesis, we previously showed that phosphorylation of wild-type and mutant HTTex1 at S13 and S16 stabilized the monomeric state and inhibited the aggregation of mutant HTTex1 *in vitro* (Deguire et al., 2018).

It was previously shown that mutant HTT from HD mouse model-derived cells shows less phosphorylation at S13/S16 than wild-type HTT (Atwal et al., 2011), and that pS13 HTT levels in a mouse striatal cell HD model as well as in HD knock-in mice are lower relative to wild type controls (Cristina Cariulo, 2019). To assess the role of phosphorylation in HTT pathology induction, we examined S13 phosphorylation levels in the brain in the zQ175 mouse model (Menalled et al., 2012). We extracted the detergent (Triton X-100)-soluble and insoluble protein fractions from 6-month-old homozygous zQ175 and heterozygous HD mice along with wild-type control mice and subjected these samples to filter retardation assays and WB. We observed very low levels of soluble HTT in homozygous zQ175 mice, and there were no significant differences in the levels of soluble or pS13 HTT between control and heterozygous mice, at least as detected by WB (Supplemental Fig. 4H). Analysis of the insoluble protein fractions revealed high levels of HTT in homozygous zQ175 mouse samples. Consistent with our previous observations, the mutant HTT proteins in the insoluble protein fraction were devoid of any pS13-phosphorylated HTT (Supplemental Fig. 4I). Together, these results combined with the previous observations that phosphorylation within N17 (pT3, pS13, pS16) inhibited mutant HTTex1 aggregation (Cariulo et al., 2017; Chiki et al., 2017; Deguire et al., 2018), indicate that reduced phosphorylation at these residues may facilitate HTT aggregation. In summary, our findings show that the TBK1-mediated reduction in HTT levels and aggregation is mediated by its ability to promote the clearance or prevent the aggregation of monomeric or soluble HTT species, rather than by the phosphorylation and disaggregation of HTT fibrils or inclusions.

**Figure 4:**
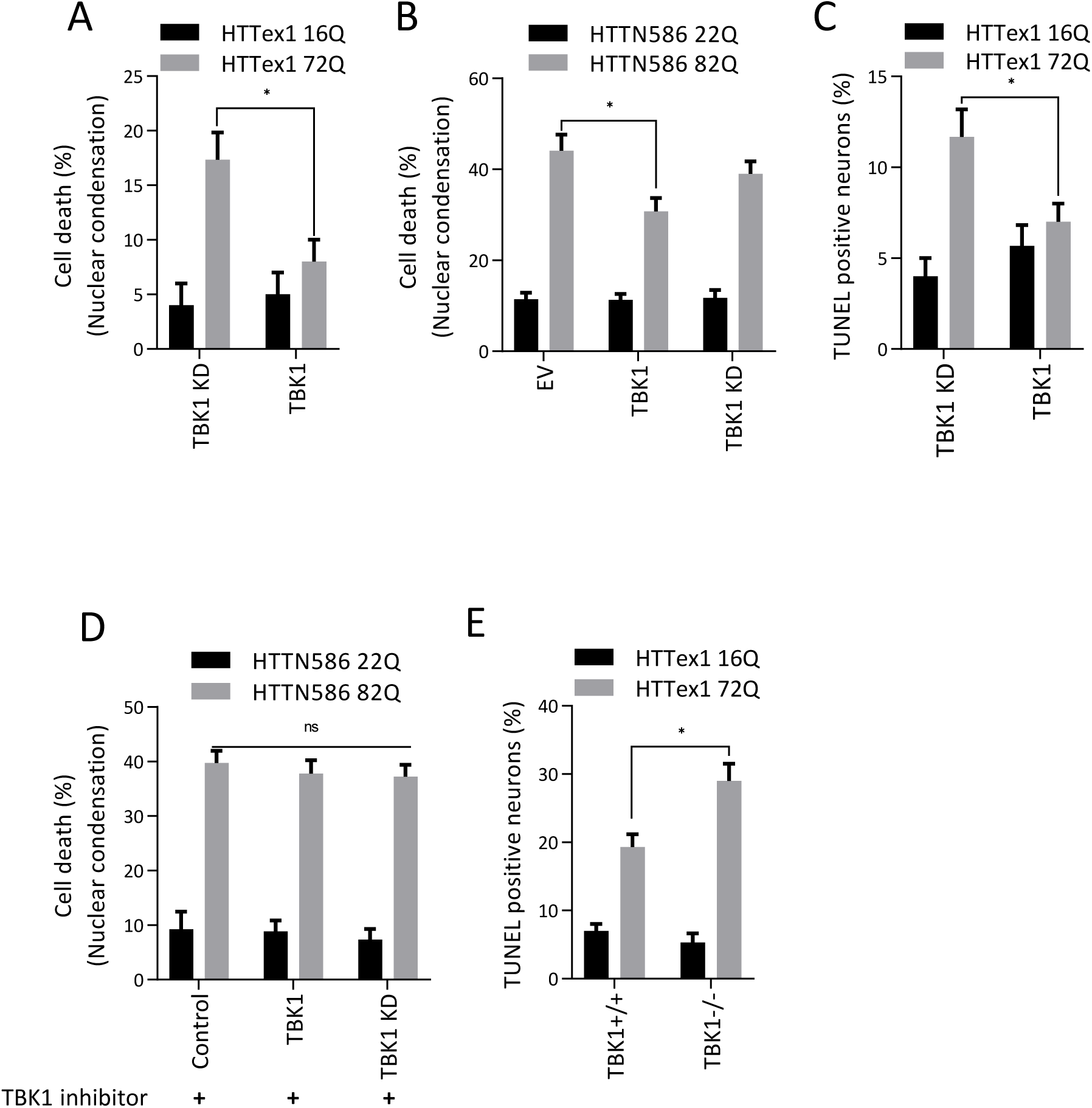
TBK1 expression leads to suppression of cytotoxic effects: **(A, B)** Quantification of cell death by a nuclear condensation assay, cells were fixed and the nuclei were stained with Hoechst dye, cells with a nuclear intensity higher than the average intensity plus two standard deviations are considered dead. (A) The percentage of cell death at DIV14, after co transfection of indicated kinases and HTTex1 plasmid in rat primary striatal neurons at DIV9, (**B**) The percentage of cell death at DIV7, after co transfection of indicated kinases and HTT N586 plasmid in mouse primary striatal neurons at DIV5. **(C)** Quantification of cell death by a TUNEL assay in rat primary striatal neurons transfected with the indicated kinases at DIV9 with the indicated HTT plasmid. The cells were fixed at DIV14, and the percentage of TUNEL-positive neurons was counted and plotted. **(D)** Quantification of cell death by a nuclear condensation assay in mouse primary cortical neurons co-transfected with the indicated HTT N586 and kinases at DIV5. After 48 Cells were treated with TBK1 inhibitor MRT 68601 for 48 hours and fixed, and the nuclei were stained with Hoechst dye. (**E**) Quantification of cell death by a TUNEL assay in mouse primary striatal neurons transfected with the indicated kinases at DIV9 with the indicated HTT plasmid. The cells were fixed on DIV14, and the percentage of TUNEL-positive neurons was counted and plotted.

The expression of mutant HTT has been shown to induce cytotoxicity in neuronal models of HD (Arbez et al., 2017; Atwal et al., 2011). Having shown that TBK1 overexpression reduced mutant HTTex1 aggregates in HEK293T cells, we sought to determine whether there would also be suppression of mutant HTTex1 cytotoxicity. We used a previously established cytotoxicity (cell death) assay based on the measurement of nuclear condensation (Cummings and Schnellmann, 2004) during neuronal cell death induced by the expression of mutant HTT (Arbez et al., 2017). We co-transfected neurons with TBK1 or a TBK1 KD variant along with 16Q or 72Q HTTex1 or HTTN586 22Q or 82Q. Consistent with our previous studies (Arbez et al., 2017), we found that HTTex1 72Q or HTTN586 82Q expression induced significant cell death as assessed by nuclear condensation, compared to HTTex1 16Q or HTTN586 22Q (Fig. 4A-B). TBK1 but not TBK1 KD co-expression partially rescued HTTex1 72Q and HTTN586 82Q-induced toxicity (Fig. 4A-B). We also observed a partial rescue of cell death induced by HTTex1 72Q upon TBK1 but not TBK1 KD co-expression in rat primary neuronal cultures using TUNEL assay (Fig. 4C). To confirm the neuroprotection mediated by overexpression of TBK1 and the dependence on its kinase activity, we repeated the HTTN586 22Q and 82Q experiments in the presence of TBK1 kinase inhibitors. We observed that treatment with the TBK1 kinase inhibitor MRT68601 (McIver et al., 2012) blocked the neuroprotective effects induced by overexpression of TBK1 (Fig. 4D). Next, we also tested whether the absence of TBK1 influences HTTex1 72Q-induced toxicity in *Tbk 1-/-, Tnf-α -/-* knockout (Hemmi et al., 2004) mouse primary striatal neurons. We observed increased toxicity in the absence of TBK1 (Fig. 4E). These findings suggest that TBK1 expression or activation may represent a viable strategy to suppress mutant HTT-induced toxicity, whereas TBK1 inhibition may further exacerbate HTT toxicity.

### TBK1 overexpression modulates HD pathology in *C. elegans*

Considering the ability of TBK1 overexpression to modulate HTT levels and its propensity to form cellular inclusions, we sought to determine whether overexpression of TBK1 could prevent mutant HTT-dependent aggregation and toxicity in a recently developed *C. elegans* model of HD (Lee et al., 2017). This model expresses the HTT N-terminal fragment up to residue 513 with 15 (Q15) or 128 glutamines (Q128) tagged with YFP in the muscle cells of the *C. elegans* body wall. In worms expressing HTTN513-YFP (Q15), HTT appeared soluble and diffusely localized to the cytosol, whereas HTT aggregates could be readily detected in the cytosol of the worms expressing mutant HTTN513-YFP (Q128) (Fig. 5A). Using these *in vivo* models, we investigated the effect of TBK1 on mutant HTTN513 aggregation and toxicity. First, we assessed whether the N17 of HTTN513 was endogenously phosphorylated in *C. elegans*. To this end, we performed a biochemical analysis of protein lysates from this model with phospho-specific antibodies against HTT S13 and T3. We observed that worms expressing mutant HTTN513-YFP (Q128) showed less phosphorylation at T3 and S13 than worms expressing wild-type HTTN513-YFP (Q15) (Fig. 5B-C), consistent with the previous reports in animal models of HD and patient-derived cells (Aiken et al., 2009; Atwal et al., 2011; Cariulo et al., 2017; Cristina Cariulo, 2019). Importantly, the detection of pT3 and pS13 HTT in *C. elegans* clearly indicates that this organism expresses kinases capable of phosphorylating these HTT epitopes.

**Figure 5:**
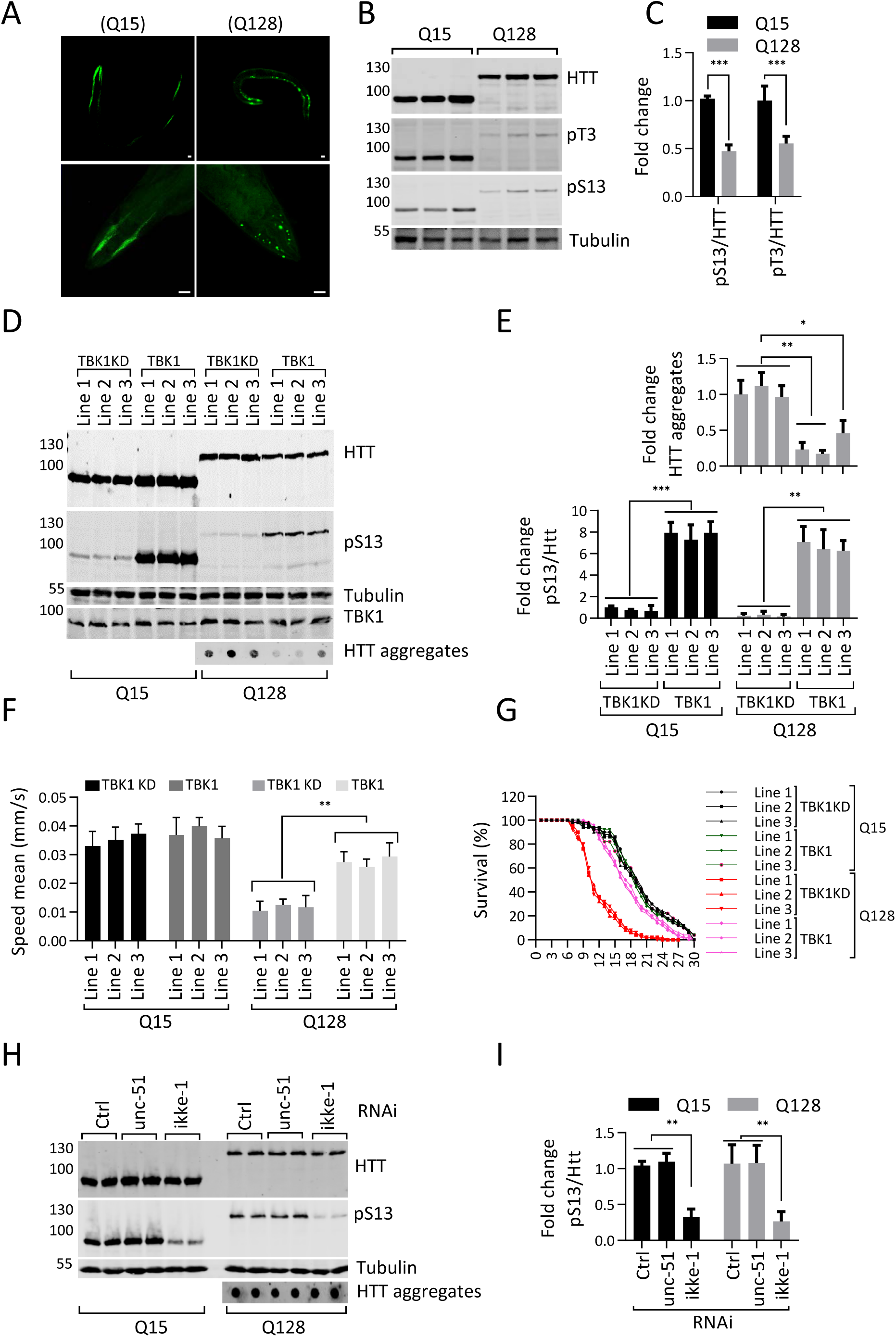
TBK1 overexpression modulates HD pathology in *C. elegans*: **(A)** Immunofluorescence image of *C. elegans* overexpressing HTT N513 Q15 and HTT N513 Q128 (scale bar 20μM). **(B)** Western blot showing HTT and pS13 and pT3 HTT levels in *C. elegans* overexpressing HTT N513 Q15 and HTT N513 Q128. **(C)** Quantification of the fold changes in ratios of HTT pS13 and pT3 to total HTT in *C. elegans* overexpressing HTT N513 Q15 and HTT N513 Q128 compared to HTT N513 Q15. **(D)** Representative western blots showing the HTT and pS13 HTT levels in three different transgenic *C. elegans* lines overexpressing TBK1 or the TBK1 KD. **(E)** The bottom graph indicates the fold change in the HTT pS13 ratios to total HTT compared to those in the kinase dead mutant line 1 from the experiments like in D. The top graph indicates the fold change in HTT aggregates compared to that in the kinase dead mutant from A. **(F)** Mobility of the Q15 and Q128 *C. elegans* lines upon overexpression of TBK1 or TBK1 KD. (**G**) Survival graph of the Q15 and Q128 *C. elegans* lines upon overexpression of TBK1 or TBK1 KD. (**H**) Representative western blots showing HTT and pS13 HTT in TBK1 orthologue (ikke-1 or unc-51) RNAi-treated *C. elegans* lines expressing HTT N513 Q15 and HTT N513 Q128. (**I**) Quantification of the fold change in the ratio of pS13 HTT to total HTT compared to non-targeting RNAi from the experiments like in H.

Approximately 81% of human kinases have orthologues in *C. elegans* (Lehmann et al., 2013; Manning, 2005). It was previously shown that transgenic overexpression of human kinases in *C. elegans* could be used to determine their effects on pathogenic disease mechanisms (Branco-Santos et al., 2017; Miyasaka et al., 2005). Therefore, we sought to determine the effect of the expression of human TBK1 in the muscle cells of the *C. elegans* HD model on HTTN513 phosphorylation, protein levels and aggregation. We generated three different transgenic lines of HTTN513-YFP (Q15) and HTTN513-YFP (Q128) worms overexpressing TBK1 or TBK1 KD (5 to 15-fold) and quantified HTT levels, phosphorylation at S13, aggregate load and worm motility, a functional phenotype affected by polyQ expansion in HTT (Lee et al., 2017). As shown in Figure 5D and E, we observed that all three lines of worms overexpressing TBK1 but not TBK1 KD showed increased phosphorylation of HTTN513-YFP at S13 in both the Q15 and Q128 models, with significantly reduced aggregate formation in the three different transgenic lines of the HTT513-YFP (Q128) model.

It was previously shown that either expression of HTTex1 or HTTN513 with expanded polyQ in *C. elegans* muscle cells in the body wall leads to aggregation-related cytotoxicity, resulting in motility defects (Lee et al., 2017; Wang et al., 2006). Moreover, the HTTN513-YFP (Q128) *C. elegans* model was shown to exhibit sustained mutant HTT-mediated toxicity and motility defects irrespective of the age of the worms (Lee et al., 2017). Therefore, we determined the biological relevance of the TBK1 overexpression-induced reduction in HTT on the modulation of polyQ-induced protein toxicity in this model. We examined the motility of HTTN513-YFP Q15 and Q128 worms co-expressing TBK1 or TBK1 KD in three different transgenic lines. As shown in Figure 5F, there was no significant difference in motility between worms co-expressing TBK1 or TBK1 KD with HTTN513-YFP (Q15). In contrast, the motility defect observed in worms expressing HTTN513-YFP (Q128) was significantly rescued upon co-expression of TBK1 but not by TBK1 KD mutant (Fig. 5F). These results suggest that TBK1 co-expression suppresses mutant HTT-induced toxicity.

Overexpression of HTTN513 Q128 within muscle cells in the body wall of *C. elegans* was shown to impair muscle function and shorten lifespan (Lee et al., 2017). Therefore, we examined the lifespan of HTT N513 Q15 and Q128 worms co-expressing TBK1 or TBK1 KD in three different transgenic lines. As shown in Figure 5G, all three lines co-expressing TBK1 KD with HTTN513-YFP (Q128) displayed a shortened lifespan (mean lifespan of ∼10 days) compared to that of three different lines co-expressing TBK1 KD with HTTN513-YFP (Q15) (mean lifespan of ∼19 days). We observed a rescue of the lifespan in all three HTTN513-YFP (Q128) lines co-expressing TBK1 (mean lifespan of ∼17 days) (Fig. 5G). These results show that TBK1 overexpression led to a significant reduction in mutant HTT aggregation and rescued mutant HTT-induced toxicity in *C. elegans*.

Next, we asked whether worm orthologues of TBK1 were involved in phosphorylating HTT at residue S13 in worms. Therefore, we tested the effect of downregulating the activity of the worm orthologues of TBK1 on HTT phosphorylation and aggregation. The worms were fed siRNAs targeting the TBK1 orthologues ikke-1 and unc-51 for 5-6 days. RT–PCR analysis of RNA from treated animals showed an efficient knockdown of their target kinases (Supplemental Fig. 5). We observed that knockdown of ikke-1 decreased pS13 levels in HTTN513-YFP (Q15) and HTTN513 -YFP (Q128) worms (Fig. 5H-I) but had no significant effect on the levels of total HTT or HTT aggregates in the Q128-expressing model during the time of the experiment. Therefore, at least one of the C-elegans TBK1 orthologues, namely ikke-1, contributes to pS13 HTT levels in these transgenic animals, thus further validating TBK1 as one of the kinases regulating phosphorylation at S13 in HTT under an endogenous condition *in vivo*.

### TBK1-induced reduction of HTTex1 72Q aggregates was dependent on autophagy

Our observation that TBK1 overexpression resulted in reduction of soluble HTTex1 and mutant HTTex1 aggregates independent of its ability to phosphorylate mutant HTTex1 suggests that TBK1 plays a role in regulating the degradation of soluble HTT. TBK1 was known to play key roles in autophagy, including the phosphorylation of several autophagy adaptors, which enhances their ability to engage LC3-II and ubiquitinated cargo (Ahmad et al., 2016; Kiriyama and Nochi, 2015; Richter et al., 2016; Weidberg et al., 2011) and to promote autophagosome maturation (Oakes et al., 2017; Pilli et al., 2012). To determine whether the TBK1-mediated reduction in HTT was mediated by proteasome- or autophagy-related mechanisms, we assessed the effects of proteasome and autophagy inhibitors on HTT levels and aggregation in HEK293T cells co-expressing HTTex1 72 Q with TBK1 or a TBK1 KD mutant. As shown in Figure 6A and B, we observed that autophagy/lysosome inhibitors but, importantly, not proteasome inhibitors blocked the TBK1-mediated reduction of both HTTex1 72Q aggregates and soluble form (Supplemental Fig. 6A).

**Figure 6:**
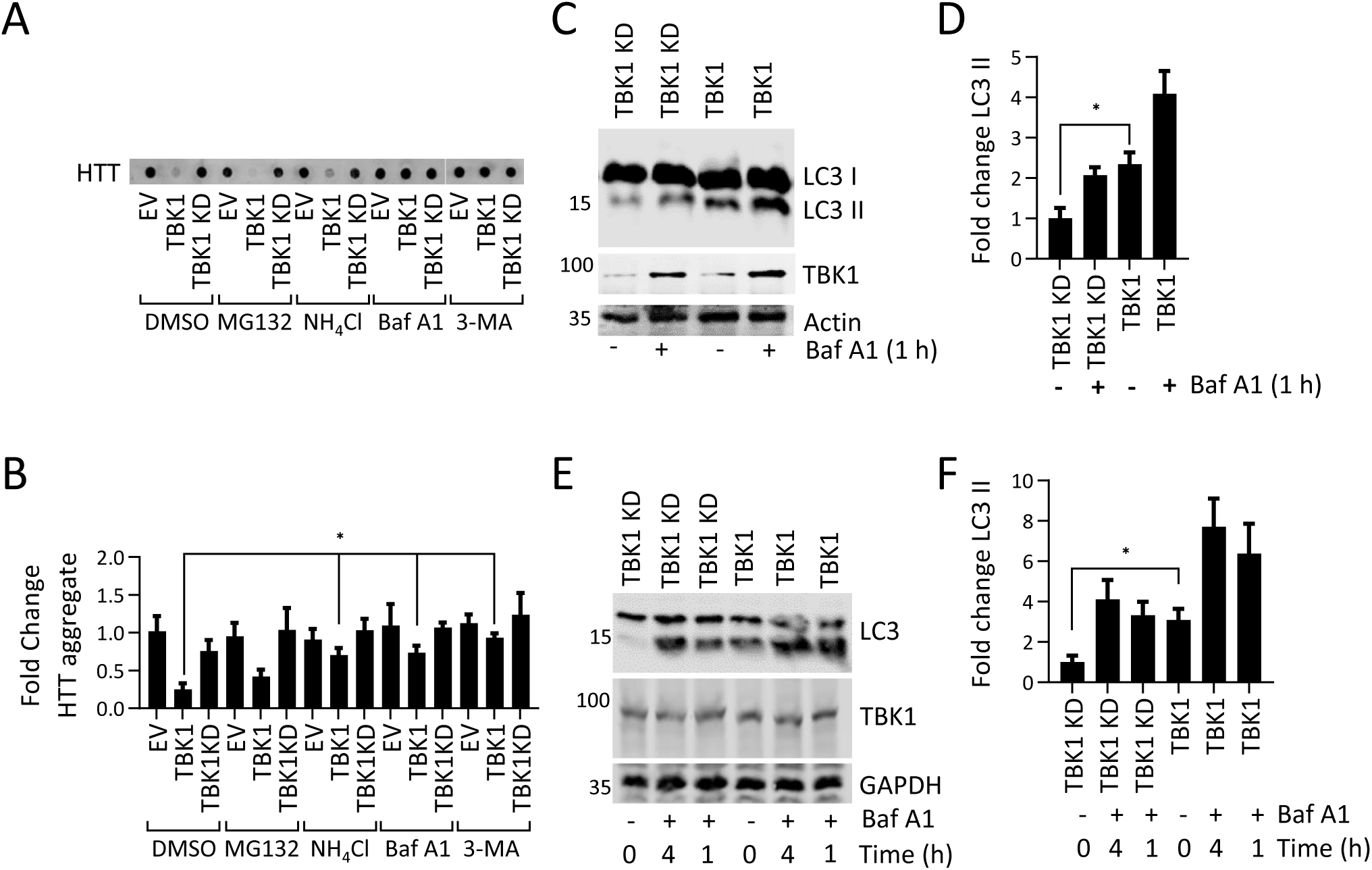
Autophagy inhibition blocks the HTTex1 72Q aggregate-reducing effect of TBK1: **(A)** Representative immunoblot of a filter retardation of HTT from the insoluble cellular fraction; HEK293T cells co-expressed HTTex1 72Q eGFP and TBK1 or TBK1 KD for 48 hrs, and for the last 16 hrs, they were treated with the indicated proteasome inhibitor (MG132-5 µM) or autophagy inhibitor (Baf A1-200 nM, NH_4_Cl-10 mM, 3-MA-5 mM). (**B**) Quantification of the fold change in HTT aggregates compared to TBK1 KD treated with DMSO from the blots in A. **(C**) Immunoblot of LC3 from soluble HEK293 cellular fractions overexpressing TBK1 or TBK1 KD for 24 hrs; for the last hour, Baf A1 (500 nM) was added as indicated. (**D**) Fold change in LC3-II levels (lower band) relative to the respective untreated kinase dead mutant normalized to actin from the blots like in F. **(E)** Immunoblot of LC3 from soluble rat primary neuronal cells overexpressing lentivirus-mediated TBK1 and TBK1 KD for 96 hours. For the last 1 or 4 hrs, Baf A1 (500 nM) was added as indicated. **(F)** Quantification of the fold change in LC3-II levels (lower band) compared to the kinase dead mutant, which was untreated, normalized to GAPDH like in E.

To determine whether TBK1 overexpression could increase autophagy in HEK293T cells and primary neuronal cells, we overexpressed TBK1 or TBK1 KD in these cellular models and assessed changes in autophagic flux by monitoring changes in the levels of LC3B (Klionsky et al., 2016) in the cell lysates. We observed that TBK1 expression increased LC3B-II levels compared to TBK1 KD (Fig. 6C-D). Bafilomycin A1 treatment resulted in a further increase in LC3B-II levels, consistent with this kinase’s ability to increase autophagy (Fig. 6C-D). An increase in autophagic flux is usually accompanied by increased clearance of the autophagy receptor p62 (Klionsky et al., 2016). Hence, we assessed p62 levels in TBK1 and TBK1 KD-expressing HEK293T cells and observed a significant decrease in p62 levels upon TBK1 overexpression compared to TBK 1KD (Supplemental Fig. 6B-C). In primary striatal neuronal cultures, the expression of TBK1, but not TBK1 KD resulted in a significant increase in LC3B-II levels (Fig. 6E-F), with an associated decrease in p62 levels (Supplemental Fig. 6D-E), indicating that TBK1 overexpression could increase autophagy in neuronal cells in a catalytic activity-dependent manner.

TBK1 was reported to phosphorylate and regulate the autophagy receptors Optineurin (OPTN) and p62 (SQSTM1), Ubiquilin-2 and the autophagy regulator RAB8B (Richter et al., 2016). Once phosphorylated, autophagy adaptors bind to ubiquitinated substrates and target these substrates for autophagy (Oakes et al., 2017; Richter et al., 2016). To validate the observed TBK1-mediated autophagy-inducing effects, we assessed whether HTT, TBK1 and autophagy adaptors co-localize in primary neurons by immunocytochemistry. We observed that endogenous, active TBK1 (phospho-serine 172) (Richter et al., 2016), co-localized with HTT, LC3B, and ubiquitin in a subpopulation of punctate structures (Supplemental Fig. 6F-G). We also observed that active TBK1 co-localized with the autophagy adaptor p62 in a few puncta in the cytosol (Supplemental Fig. 6H-I). In summary, our data suggest that TBK1 expression promotes the general cellular clearance mechanism of soluble HTT and prevents its accumulation and aggregation by enhancing autophagy.

### Upon disruption of TBK1 interactions with autophagy adaptor proteins, its effects on HTT levels and aggregate formation become strongly dependent on the phosphorylation state of HTT

To uncouple the effects of phosphorylation-dependent changes and TBK1-induced enhancement of autophagy on HTT turnover and aggregation, we investigated the effect of expressing a characterized TBK1 mutant incapable of binding to autophagy adaptors (lacking amino acids 690 to 713, within the coiled-coil domain (Oakes et al., 2017)). It was also observed that TBK1 with a deletion of amino acids 690 to 713 (Δ690-713) could still phosphorylate known substrates, such as IRF3 (Freischmidt et al., 2017; Freischmidt et al., 2015). To further dissect the underlying mechanisms, we examined the effects of the expression of TBK1 Δ690-713, TBK1-wild type or TBK1 KD on the levels and aggregation of phosphorylation-competent HTTex1 72Q and phosphorylation-deficient S13A/S16A mutant in HEK293 cells. Figure 7A and B show that both TBK1 Δ690-713 and TBK1-wild type could phosphorylate HTTex1 72Q resulting in a significant reduction in HTTex1 72Q aggregates. However, blocking phosphorylation at S13 and S16 (using S13A/S16A mutations) blocked the ability of TBK1 Δ690-713 to reduce HTTex1 72Q aggregation (Fig. 7A-D). Unlike wild type-TBK1, co-expression of TBK1 Δ690-713 had minimal effects on the levels of soluble HTT. These results indicate that the TBK1-mediated reduction of mutant HTT aggregates was dependent on its autophagy adaptor-binding function and that this mechanism was the primary contributor to the phosphorylation-independent reduction of aggregates of HTT S13A/S16A upon co-expression of TBK1. Upon removal of the 690-713 region, TBK1 did not phosphorylate p62 or OPTN, did not activate autophagic flux (Supplemental Fig. 6J) and its effect on HTT aggregation became dependent on the phosphorylation state of HTT at S13/S16. Importantly, these data highlight that phosphorylation at S13 was sufficient to prevent mutant HTT aggregation without requiring the TBK1-mediated enhancement of autophagy or the mutant HTTex1-lowering effect. These results indicate that phosphorylation at S13/S16 may suppress mutant HTT aggregation, consistent with our recent *in vitro* studies (Deguire et al., 2018) and phosphomimetic studies using a mouse model (Gu et al., 2009) and cell lines (Atwal et al., 2011; Branco-Santos et al., 2017; Thompson et al., 2009).

**Figure 7:**
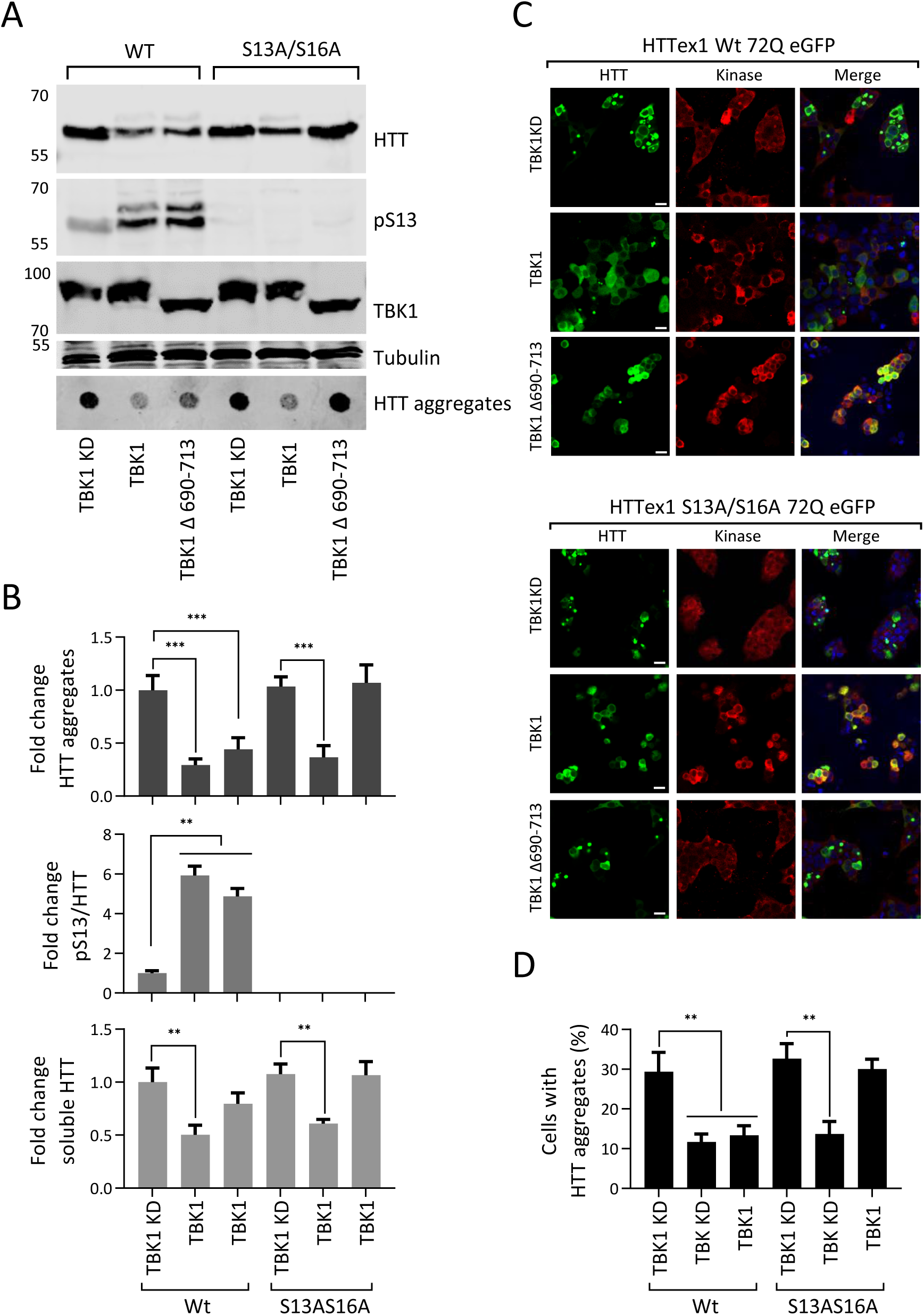
TBK1 deficient in the autophagy adaptor domain does not reduce 72Q eGFP S13A/S16A aggregates: Western blot of soluble HTT, HTT pS13 and aggregates upon co-expression of the HTTex1 72Q eGFP or HTTex1 72Q eGFP S13A/S16A variant with TBK1 KD, TBK1 or TBK1 Δ690-713 for 48 hours in HEK293 cells. **(B)** The graph indicates the calculated fold changes in HTT, HTT pS13 ratio to total HTT and HTT aggregates all compared to the levels in TBK1 KD from the experiments in A (bottom panel normalized to Tubulin). (**C**) Representative immunofluorescence images of co-expression TBK1 KD, TBK1 or TBK1 Δ690-713 with HTTex1 72Q eGFP or its phosphorylation-deficient variant S13A/S16A for 48 hours in HEK293 cells (scale bar 20μM). (**D**) The graph indicates the percentage of co-transfected cells presenting aggregates upon expression of the indicated kinase, from the experiments like in C.

## Discussion

Increasing evidence from *in vitro*, cellular and animal model studies shows that phosphorylation or phosphorylation-mimetics at S13 and/or S16 attenuates mutant HTT aggregation and protects against mutant HTT (full-length or N-terminal fragment)-induced toxicity (Arbez et al., 2017; Atwal et al., 2011; Branco-Santos et al., 2017; Deguire et al., 2018; Thompson et al., 2009). However, whether inhibiting mutant aggregation is the primary mechanism underlying the protective effects of phosphorylation at these residues remains unknown. This is in part because the majority of previous studies aimed at dissecting the role of S13/S16 phosphorylation in cellular and animal models were based primarily on approaches relying on the use of mutations to mimic or block phosphorylation, rather than *bona fide* phosphorylation and thus did not account for the dynamic nature of phosphorylation. To address this knowledge gap and elucidate the role of phosphorylation at S13 and S16 in HTT aggregation, turnover and toxicity we set out to identify kinases capable of increasing S13 HTT phosphorylation and identified TBK1 as a kinase that efficiently phosphorylates wild-type and mutant HTT at S13 and S16 *in vitro* and S13 in cellular and animal models of HD.

Herein, we showed that TBK1 co-incubation with wild-type or mutant HTTex1 or longer N-terminal HTT fragments resulted in phosphorylation of both S13 and S16 (Fig. 1C and Supplemental Fig. 1A-B). In contrast to IKKβ, which was shown previously to preferentially phosphorylate wild-type HTTex1 at residue S13 (Thompson et al., 2009), TBK1 phosphorylated both wild-type and mutant HTT *in vitro*. In our hands, IKKβ phosphorylated both S13 and S16, although much less efficiently than TBK1 (Supplemental Fig. 2A-B). Furthermore, TBK1 co-expression/ overexpression robustly and efficiently increased S13 phosphorylation of both wild-type and mutant HTT in cells by approximately 5-to 10-fold compared to 3 to 4-fold by IKKβ (Fig. 1D-E) and lead to stronger HTT-lowering effects, especially for mutant HTTex1. To the best of our knowledge, this is the first report demonstrating that HTT is a substrate of TBK1. The discovery of TBK1 provides an efficient tool for manipulating HTT phosphorylation levels in cells and enabled us to systematically determine how modulating phosphorylation at these residues influenced HTT aggregation (Fig. 8A), clearance and toxicity in different models of HD.

**Figure 8:**
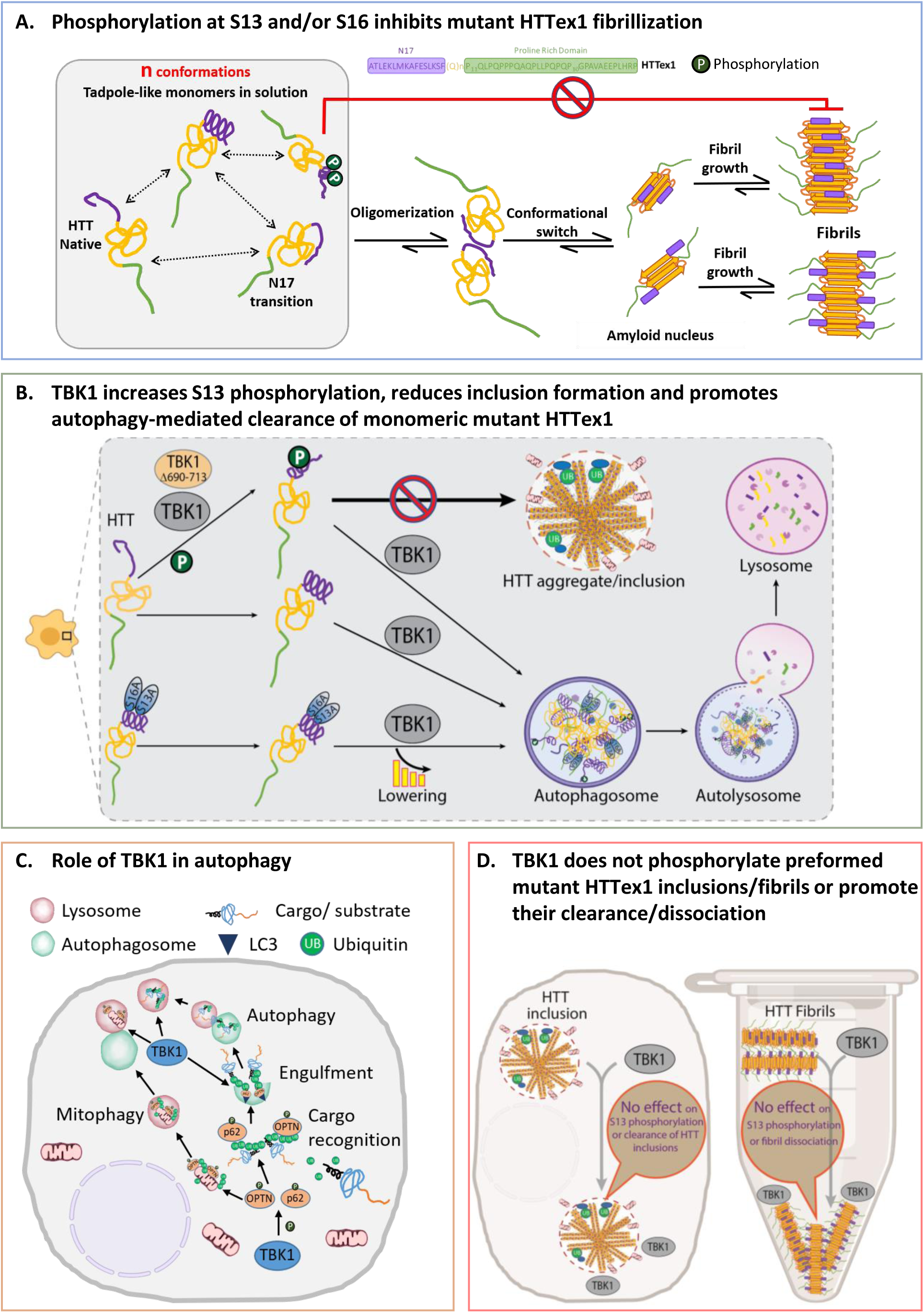
Schema of the effect of HTT phosphorylation and TBK1 on aggregation: (**A**) Effect of S13 and/or S16 phosphorylation on HTTex1 aggregation. HTTex1 forms a tadpole-like monomer structure and N17-driven interaction may further drive HTTex1 oligomerization and fibril formation. Phosphorylation at S13 and S16 destabilizes the helix at N17 that leads to inhibition of formation of HTTex1 fibrils. (**B**) Effect of TBK1 on HTT in the cellular HD model. TBK1 and TBK1 Δ690-713 both phosphorylate monomeric or soluble HTTex1 72Q at S13 residue leading the reduction of the aggregation, but TBK1 also reduces the aggregation of both HTTex1 and HTTex1 S13A/S16A through the lowering of the HTT levels at monomeric stage by enhancing the autophagic degradation. TBK1 Δ690-713 only reduces the aggregation of phosphorylation-competent HTTex1 but do not reduces the aggregation of HTTex1 S13A/S16A highlighting the role of S13 phosphorylation in aggregation inhibition. (**C**) TBK1 effects on different stages of autophagy. TBK1 is known to phosphorylate autophagy adaptors p62, OPTN that bind ubiquitinated substrates like proteins or damaged mitochondria and recruit LC3 and increase the autophagic clearance of substrates. (**D**) TBK1 does not phosphorylate HTTex1 in inclusion/ fibrils or promote their clearance/ disaggregation. When HTT was already aggregated in cells or when in fibril form *in vitro*, possibly S13 residue is not accessible for TBK1 phosphorylation as N17 was shown to be collapsed on fibrillar core or it is near the core. Also, TBK1 mediated autophagy flux was not able to induce the lowering of HTTex1 inclusion. There was no disaggregation of fibrils up on TBK1 kinase *in vitro*.

### TBK1 regulates the autophagy-mediated clearance of monomeric HTT

Our results demonstrate that TBK1 modifies HTT aggregation and toxicity through autophagy-mediated regulation of HTT clearance. TBK1 overexpression in cells reduced the levels of soluble HTTex1 and prevented the accumulation of mutant HTTex1 inclusions in a kinase activity-dependent manner, whereas TBK1 knockdown, knockout or inhibition resulted in increased HTT levels. These observations suggest that TBK1 regulates mutant HTT levels and inclusion formation by activating clearance mechanisms such as proteasome- or autophagy-mediated protein clearance.

TBK1 levels and activity have been implicated in regulating many aspects of autophagy. For example, *TBK1* gene duplication in human iPSC-derived retinal cells or overexpression of TBK1 in RAW264.7 cells leads to increased LC3II and increased autophagy (Pilli et al., 2012; Ritch et al., 2014). TBK1-mediated phosphorylation of the autophagy adaptor proteins OPTN and p62 represents a rate-limiting step in the autophagic degradation of misfolded protein aggregates. Several studies have also shown that TBK1 acts as a scaffolding protein and plays an important role in regulating autophagic activity by regulating protein targeting to autophagosomes, autophagosome formation, autophagosome engulfment of substrates (Korac et al., 2013; Matsumoto et al., 2011), and fusion of autophagosomes to autolysosomes (Pilli et al., 2012). TBK1 depletion negatively impacts all of these processes and causes decreased autophagic activity (Oakes et al., 2017). Defects in autophagy and autophagosome formation and clearance have been implicated in the pathogenesis of neurodegenerative diseases (Nixon, 2013; Scrivo et al., 2018).

Consistent with its role in autophagy (Oakes et al., 2017), we observed that TBK1 co-localized with mutant HTTex1 inclusions formed by overexpression of both proteins in HEK293 cells and rat striatal primary neurons (Supplemental Fig. 7A-C). TBK1 overexpression in our HTT aggregation model led to an increase in autophagic flux and the phosphorylation of the autophagy adaptors OPTN and p62 (Supplemental Fig. 6J). To determine whether TBK1-mediated enhanced clearance or reduced aggregation of HTT was due to its interaction with and/or phosphorylation of HTT monomers, aggregates or both, we assessed the effect of TBK1 overexpression on HTT levels and aggregation in different HD models. Interestingly, we observed that the TBK1-mediated reduction in HTT levels and aggregation were strongly dependent on TBK1 kinase activity and its ability to bind to the autophagy adaptor OPTN, but occurred independently of phosphorylation at S13 (Fig. 8B). However, upon disruption of TBK1 interactions with the autophagy adaptors OPTN and p62, the effects of TBK1 on HTT levels and aggregation became strongly dependent on both its kinase activity and HTT phosphorylation at S13. These findings suggest that the primary mechanism underlying the beneficial effects of TBK1 is mediated by its ability to enhance autophagy, possibly through phosphorylation of the autophagy adaptors OPTN and p62 (Fig. 8C). Preliminary studies from our group (Maharjan et, al unpublished) investigating the PTM-dependent interactome of soluble monomeric HTT showed that phosphorylation at S13 and S16 promoted interactions between mutant HTT and many autophagy, folding and endocytic pathway components, suggesting that phosphorylation at these residues could also regulate HTT levels by promoting its interactions with key mediators of cellular proteostasis mechanisms. The proposed role of HTT as an autophagy scaffold protein (Ochaba et al., 2014) is also consistent with this hypothesis. Previous study showed that phosphorylation of mutant HTT at S13/S16 significantly destabilized the α-helix and reduced the rate and extent of aggregation *in vitro* (Deguire et al., 2018). Our results further support the hypothesis that enhancing phosphorylation at S13 is sufficient to induce a significant reduction in HTT aggregation and inclusion formation.

Our observation that TBK1 was not able to induce the clearance of HTT post-aggregation is consistent with previous studies showing that autophagy induction by rapamycin treatment had no effect on the levels of HTT aggregates once mature aggregates were formed but reduced HTT aggregate formation and cell death when applied at the earlier stages of the aggregation process (Ravikumar et al., 2002). Interestingly, TBK1 was not capable of phosphorylating mutant HTTex1 fibrils *in vitro*, HTT aggregates in cells or inducing the clearance of mutant HTT aggregates once they were formed (Fig. 8D). Together, these findings suggest that the beneficial effects of TBK1 are mediated by its activity-dependent regulation of the phosphorylation and degradation of monomeric, but not aggregated forms of HTT.

### Therapeutic Implications

The discovery of TBK1 as a kinase that efficiently phosphorylated HTT at S13 presents unique opportunities to directly assess the phosphorylation at these residues or selective autophagic degradation of monomeric HTT as therapeutic targets for the treatment of HD. Several lines of evidence support these hypotheses: 1) small molecules such as GM1 (Di Pardo et al., 2012) and N6-furfuryladenine (Bowie et al., 2018) were shown to increase S13/S16 phosphorylation and restore normal motor behavior in the YAC128 HD mouse model; 2) a BACHD mouse model expressing S13D/S16D HTT showed a rescue of symptoms of HD pathology, such as motor deficits and reduced aggregate formation (Gu et al., 2009); and 3) our results in *C. elegans* showed that TBK1-mediated increased phosphorylation at S13 resulted in a significant reduction of HTT aggregates and improved worm lifespan and motility. Furthermore, we showed that increasing TBK1 levels or activity resulted in a robust increase of HTT phosphorylation and enhancement of autophagy, leading to reduced HTT levels and suppression of HD pathology and toxicity, suggesting that targeting TBK1 represents a viable strategy for the treatment of HD.

Increasing evidence also suggests that dysfunction in autophagy and other protein clearance mechanisms play central roles in the pathogenesis of neurodegenerative diseases, including HD (Croce and Yamamoto, 2019; Nixon, 2013) (see supplemental discussion). Herein, we showed that TBK1 expression/activation helped to overcome the autophagy defects induced by HTT misfolding and aggregation in HD models. Our data suggest that TBK1 expression/activation increases the phosphorylation of the cargo/substrate recognition adaptors OPTN and p62. Phosphorylation on OPTN or p62 was shown to facilitate the autophagy-dependent clearance of damaged/dysfunctional mitochondria (Matsumoto et al., 2015; Richter et al., 2016), a common defect in many neurodegenerative diseases (Rodolfo et al., 2018; Wang et al., 2019) and misfolded proteins and aggregates of proteins linked to neurodegenerative diseases, such as mutant SOD1 (Korac et al., 2013) and HTT (Matsumoto et al., 2011). TBK1-mediated enhancement of cargo/substrate recognition by the adaptors OPTN and p62, and *OPTN* mRNA levels were shown to be partially reduced in HD patient brain tissues (Hodges et al., 2006). In cellular models of HD (expressing HTTex1 103Q), OPTN knockdown led to an increase in HTT aggregation (Korac et al., 2013). Although the role of OPTN in the progression of HD is currently unknown, reducing OPTN activity or levels may confer susceptibility to HTT aggregation because of its role in autophagic clearance of protein aggregates. Therefore, pharmacological enhancement of the TBK1 pathway could activate the available pool of OPTN, which may, in turn, increase the recognition and clearance of HTT. Previous studies suggested that general autophagy induction in HD mouse models reduce aggregate load and improves motor performance and lifespan (Ravikumar et al., 2004; Rose et al., 2010; Tanaka et al., 2004). Our results suggest that TBK1 is a strong candidate for activating specific steps involved in the autophagy cascade (Fig. 8 D), and for the reduction of the autophagy substrates.

Although increasing TBK1 activity represents an attractive strategy for lowering the levels of soluble HTT species and preventing HTT aggregation, our findings showed that TBK1 did not act on or promote the phosphorylation or degradation of preformed HTT aggregates. However, given the dynamic nature of HTT aggregation and its reversibility, for example, under conditions where HTT expression was suppressed or lowered, it is likely that the TBK1-mediated reduction in soluble HTT could significantly contribute to a reduction of the levels of HTT aggregate formation by means of preventing the growth of existing aggregates and/or the formation of new aggregates. Our current data and previous observations by Ravikumar et al. (Ravikumar et al., 2002) suggest that early activation of autophagy could be beneficial and promote the clearance of mutant HTT aggregates in HD. One consideration is that with increasing age and at advanced disease stages, autophagy-lysosome and proteasome abnormalities become more apparent in HD (Cortes and La Spada, 2014; Seo et al., 2004). However, given that patients with HD can be identified at pre-symptomatic disease stages through genetic testing, therapeutic autophagy modulation alone or in combination with HTT-lowering strategies before disease onset is theoretically possible in HD and might be necessary for a significant beneficial effect.

### Implications for other neurodegenerative diseases

Several stages of autophagy, such as substrate sequestration and autophagosome formation, autophagosome-lysosome-endosome fusion, and lysosomal digestion, were shown to be impaired in ALS, AD, PD and FTD (Nixon, 2013). Several mutations in TBK1 and OPTN cause ALS and were shown to impair autophagy (Oakes et al., 2017). Targeting autophagy induction in these diseases by mTOR inhibitors, AMPK activation or selective autophagy induction by overexpression of specific genes has been shown to be beneficial in many different cellular and mouse models of neurodegenerative diseases such as ALS, AD, PD and FTD (Menzies et al., 2017; Oueslati et al., 2013; Qi et al., 2012; Scrivo et al., 2018; Sorrentino et al., 2017; Spilman et al., 2010; Wu et al., 2011). Considering the role of TBK1 in autophagy and the increasing evidence implicating its activity in the clearance of misfolded protein aggregates and ALS, it is plausible that pharmacological enhancement of TBK1 activity or selective targeting of TBK1-dependent autophagic pathways represent attractive and viable therapeutic strategies for the treatment of multiple neurodegenerative diseases, provided that TBK1 levels are not severely reduced in the disease. Indeed, we do not observe any changes in TBK1 and TBK1-pS172 in the brains of R6/2 transgenic HD mouse model and HD postmortem brain tissue (Supplemental Fig. 7D-I). Recent success stories in finding kinase activators for Lyn, PKCδ and AMPK kinase (Bessa et al., 2018; Cool et al., 2006) in the fields of diabetes drug development suggest that finding activators of TBK1 activity is feasible and may provide a therapeutic option to modulate HD and other neurodegenerative diseases.

## Acknowledegments

This work was supported by funding from CHDI and EPFL and NINDS (R01 NS086452 and R21 NS083365). We are grateful to Anne-Laure Mahul-Mellier, Galina Limorenko, Johannes Burtscher, Niran Maharjan, Senthil Kumar Thangaraj and Somanath Jagannath, EPFL for critical review of the manuscript. We thank Rajasekhar Kolla for preparing the Graphical abstract. We are grateful to staff at the Bio-imaging Core Facility (BioP, EPFL) for their technical support. We thank Driss Boudeffa, EPFL for the production of lentiviral vectors; Jonathan Ricci, EPFL for assisting *in vitro* kinase validation; Elena Gasparotto, Lorène Aeschbach, EPFL for assisting with neuronal cell culture experiments. We thank Naveed Ziari and Kevin S. Hof, EPFL-LISP-IBI-SV for assisting with experiments in Fig. 5 B, C, H, I and Supplemental Fig. 5 and initial help with the setting up of *C. elegans* model. We thank the proteomics core facility at EPFL’s School of Life Sciences and Functional Genomics Center Zurich (FGCZ), ETH Zurich for their support with the mass spectrometry analysis. We are grateful to Prof. Elise A. Kikis, The University of the South, Sewanee, Tennessee, for the kind gift of HD *C. elegans* model and Prof. Shizuo Akira, WPI Immunology Frontier Research Center, Osaka University, Japan for the kind gift of *Tbk1-/-* mouse model used in this study.

## Contributions

HAL, RNH and AC conceived the project; HAL, RNH and AC designed the experiments; RNH, performed the experiments in Figs. 2 A-B and G, H; Figs. 3, 4 A, C, E; Figs. 5, 6, 7; Supplemental Figs. 3, 4 A-F and H, I and Supplemental Figs. 5, 6, 7 A-C, E, F. AC performed the experiments in Fig. 1 A-C; Supplemental Figs. 1, 2, and 4 G; NZ and KH and LM assisted RH for the experiments in Fig. 5 B, C, H, I and Supplemental Fig. 5. LM and JA supervised the NZ and KH. PM performed the experiments in Fig. 2 C-F. ACa and LP designed, supervised experiments in Fig. 2 C-F. NA performed the experiment in Fig. 4 B, D; CAR conceived the experiments by NA. CL performed the experiment in Supplemental Fig. 7D; GB conceived the experiments by CL. MKSB performed, analyzed and wrote materials and methods for the experiment in Supplemental Fig. 7G-I; RF and MAC supervised the ethical collection, donor interaction, clinical assessment and processing of the human tissues for Supplemental Fig. 7G-I. RF co-designed experiments in Supplemental Fig. 7G-I. RNH, AC and HAL analyzed the data, RNH, AC and HAL wrote the draft and completed the final manuscript writing with the contributions from all authors.

