## Supplemental Information for "TBK1 regulates autophagic clearance of soluble mutant huntingtin and inhibits aggregation/toxicity in different models of Huntington’s disease"

#### Results:

##### Identification and validation of TBK1 as a kinase that efficiently phosphorylates HTT at S13 and S16

To identify the kinases responsible for phosphorylating HTT at S13 and S16, approximately 300 purified serine-threonine kinases (Kinexus *in vitro* kinase and phospho-peptide testing (IKPT) services-(HTTp://www.kinexus.ca/ourServices/substrate_profiling/substrate_profiling/phosphopeptide_kinase.html) were screened using a peptide that comprised the N-terminal 17 residues of HTT (N17) and exon1 of the HTT protein (HTTex1) as a substrate (Fig. 1A). TBK1 was the only kinase that efficiently phosphorylated HTTex1 *in vitro* at S13 and S16. To validate the specificity of TBK1, we assessed its ability to phosphorylate recombinant N-terminal HTT fragments 2-140 and 1-171 produced *in vitro* in *E. coli*. HTT 2-140 and HTT 1-171 contain several other potential phosphorylation sites in the C-terminus. The extent of phosphorylation was monitored using mass spectrometry and previously validated phospho-antibodies against T3 (pT3), S13 (pS13) and S16 (pS16) (Bustamante et al., 2015; Deguire et al., 2018). Supplemental Fig. 1 A and B shows a time-dependent increase in TBK1-mediated phosphorylation of the longer N-terminal fragments HTT 2-140 and HTT 1-171, predominantly di-phosphorylation at S13 and S16, along with a small amount of tri-phosphorylated and traces of tetra-phosphorylated species. For these substrates and HTTex1 a signal was observed for only the pS13 and pS16 antibodies, demonstrating that TBK1 did not phosphorylate HTTex1 at T3 (Supplemental Fig. 1A-B lower panel). Consistent with these data, we did not observe any phosphorylation of a HTTex1 mutant in which both residues were mutated to aspartate (S13D/S16D), even after extended periods of incubation with TBK1 (Supplemental Fig. 1 C-E).

The TBK1 family kinase IKKβ was previously reported to phosphorylate HTT at S13/S16 (Thompson et al., 2009). Therefore, we sought to compare the efficiency of TBK1- and IKKβ-mediated phosphorylation of HTTex1. To this end, we compared the kinetics and efficiency of HTTex1 phosphorylation by these two kinases. As shown in Supplemental Fig. 2A-B, we observed significantly faster phosphorylation by TBK1 than by IKKβ at both S13 and S16 residues, and this effect was already apparent after only 30 mins. IKKβ seemed to require a longer time to phosphorylate HTT, as we observed phosphorylation after 6 hours but not at 3 hours. After 6 hours, TBK1 resulted in almost complete di-phosphorylation, whereas IKKβ showed predominantly mono-phosphorylated HTT even after 16 hours, with a significant amount of non-phosphorylated HTT remaining (Supplemental Fig. 1C-D). Altogether, these findings demonstrate that TBK1 mediates efficient and site-specific phosphorylation of recombinant N-terminal HTT fragments at S13 and S16 *in vitro*, consistent with its role as one of the natural kinases that regulate HTT phosphorylation and cellular properties *in vivo*.

**TBK1 interacts with HTTex1 in HEK293 cells**

Having established that TBK1 phosphorylated HTTex1 and other fragments in different cellular models, we sought to determine whether TBK1 interacted directly with its HTT substrates. To this end, we co-expressed TBK1 or TBK1 KD with wild-type or mutant HTTex1 bearing the S13A/S16A mutation or no mutation for 48 hours in HEK293 cells and immunoprecipitated HTT. We observed that TBK1 specifically coimmunoprecipitated with HTTex1 irrespective of the presence or absence of the S13A/S16A mutation (Supplemental Fig. 2C-D). These results suggest that both wild-type and mutant HTTex1 may interact with TBK1 in cells and that the phosphorylation-deficient mutations at S13 and S16 of HTT do not influence the HTT-TBK1 interaction.

**TBK1 downregulation does not affect the HTT pS13 phosphorylation**

Having established that TBK1 phosphorylated full-length HTT and various N-terminal HTT fragments and having identified and validated the phosphorylation sites, we next sought to determine whether TBK1 is one of the natural enzymes that regulated the phosphorylation of HTT in primary neurons and *in vivo*. Towards this goal, we assessed changes in the levels of pS13 HTT upon genetic knockdown or pharmacological inhibition of TBK1 in rat primary striatal neurons. Due to its known role in inflammatory disorders and cancer, several TBK1 inhibitors were developed for the treatment of related diseases (Hasan and Yan, 2016; Oakes et al., 2017; Yu et al., 2015). We treated rat primary striatal neurons with TBK1 kinase inhibitors for 48 hours or with siRNA for 72 hours and assessed the effects on HTT and HTT pS13 levels using pS13-specific antibodies. Supplemental Fig. 3 E and F show that TBK1 knockdown led to no significant change in HTT pS13 levels, but increased endogenous HTT levels by approximately 1.5-2-fold, and we observed no change in pS13 levels but there was a trend towards an increase in endogenous HTT levels in the inhibitor-treated striatal neuronal cells (Supplemental Fig. 3 G-H). While these results clearly demonstrate that TBK1 is involved in regulating the turnover of endogenous HTT, no significant change in HTT pS13 levels caused by either inhibitor treatment or siRNA treatment was observed. This might be due to sufficient residual kinase activity that could phosphorylate endogenous HTT. Alternatively, other kinases may phosphorylate HTT at S13.

To clarify whether endogenous TBK1 phosphorylated HTT at S13 in the mammalian brain, we used a mouse model of *Tbk1*-/- on the *Tnf-α* -/- background (Hemmi et al., 2004). We dissected the cortices of P0 mouse pups from the *Tbk1* +/+, *Tnf-α* +/+ and *Tbk1* -/-, *Tnf-α* -/- strains, lysed the tissues, and subjected the total protein extract to WB analysis to determine HTT and HTT pS13 levels. We used this double knockout model because *Tbk1* knockout was shown to be embryonic lethal, whereas *Tbk1* -/-, *Tnf-α* -/- double knockout animals survive without any major phenotype impairments (Perry et al., 2004). As shown in Supplemental Fig. 3I and J, consistent with knockdown and inhibitor experiments, we observed no change in pS13 HTT levels but there was an increase in the total HTT level. These observations may be explained by compensatory mechanisms involving other kinases, or homeostatic mechanisms involving downregulation of relevant phosphatases. However, the increased levels of HTT upon silencing or inhibition of TBK1 indicated a role for TBK1 in HTT clearance.

**TBK1 level is not affected in R6/2 transgenic HD mouse model and HD patients**

Having determined the relevance of TBK1 for HTT we sought to determine if the levels of TBK1 are altered in HD and HD models. We examined the levels of active TBK1 (phospho S172 =pS172) total TBK1 in R6/2 transgenic HD mouse model and human postmortem brain tissues and active TBK1 pS172 in HD patient brain tissue microarray. We observed no change TBK1 or TBK1 pS172 levels in R6/2 transgenic HD mouse brain tissue lysates at any time-points (4, 8, 12 weeks) (Supplemental Fig. 7D). We then obtained five HD and five non-HD post mortem cortical tissue and analyzed the TBK1 level in tissue lysates. We observed a trend of decrease in both the total or active TBK1 levels from these tissues, which was not statistically significant (Supplemental Fig. 7E-F). These trend of decrease in TBK1 and active TBK1 levels lead us to study the levels in a larger set of human samples. For this, we chose to use the tissue microarray approach utilizing the TBK1 and TBK1 pS172 antibody. We did not observe any significant changes in active TBK1 pS172 expression in 21 HD samples compared with 18 control samples from the middle temporal gyrus tissue microarray (total TBK1 antibody was not suitable for immunohistochemistry on paraffin tissue microarrays) (Supplemental Fig. 7G-H). Moreover, there was no apparent correlation between the expression levels of active TBK1 and HTT stained by TBK1 pS172 and HTT 1C2 antibodies, respectively (Supplemental Fig. 7I). These results suggest TBK1 levels may not be drastically changed in HD cases.

**Discussion:**

**TBK1 does not phosphorylate or reduce the already formed HTT aggregates**

Our study also demonstrated that TBK1 was not capable of phosphorylating mutant HTTex1 at S13/S16 in fibrils *in vitro* or HTT aggregates in cells. These observations suggest that the Nt-17 domain adopts a different conformation in the aggregated state of mutant HTT and/or that TBK1 does not interact with aggregated forms of HTT. Indeed, previous solid-state NMR studies on HTTex1 fibrils showed that N17 was less mobile and was engaged in interactions with the amyloid core or PRD domain in the fibrillar state (Fig. 8A) (Bugg et al., 2012; Lin et al., 2017). The inaccessibility of N17 in the fibril state suggests that post-aggregation phosphorylation or potentially other PTMs of N17 are less likely to occur (Fig. 8D). These findings imply that the Nt-17 PTM-dependent regulation of HTT aggregation and clearance is likely mediated by the aggregation-inhibitory effects of PTMs at the monomer level or their ability to selectively target monomeric HTT for degradation. Several lines of evidence support this hypothesis: 1) we failed to detect phosphorylation at S13 in insoluble HTT fractions and aggregates from cellular and animal models of HD, whereas phosphorylation at S13 was readily detectable in the soluble fractions; 2) several studies reported reduced phosphorylation of mutant HTT at Nt-17 (Aiken et al., 2009; Atwal et al., 2011; Cariulo et al., 2017); and 3) previous and ongoing studies from our group showed that the majority of the Nt-17 modifications at the monomeric level (phosphorylation at T3, S13, S16 and S13/S16; ubiquitination; SUMOylation; methionine oxidation) inhibit mutant HTTex1 aggregation (Chiki et al., 2017; Deguire et al., 2018; Steffan et al., 2004; Thompson et al., 2009). We have yet to identify an Nt-17 PTM that enhances the aggregation of mutant HTTex1 *in vitro*.

**HD models show autophagy defects**

Studies of autophagy processes in the HD human brain and mouse models showed that mutant HTT causes 1) defective autophagy cargo/substrate recognition and loading, leading to the accumulation of empty autophagosomes (Martinez-Vicente et al., 2010); 2) defective autophagosome transport near the lysosomes (Wong and Holzbaur, 2014); and 3) reduced lysosomal activity, trafficking and translocation (del Toro et al., 2009; Qi et al., 2012). More recently, HTT has also been shown to have an autophagy-inducing domain and to play a possible role in regulating autophagy (Martin, Heit et al. 2014). Bioinformatics analyses predicted 11 linear motifs known as LC3-interacting repeats (LIRs; a domain necessary for delivery to autophagosomes) within full-length HTT (Kalvari et al., 2014). HTT has also been demonstrated to function as a scaffold protein for selective macroautophagy, as well as to promote selective autophagy through its ability to interact with p62 and ULK1 simultaneously, thus modulating both cargo recognition efficiency and autophagosome formation initiation (Ochaba et al., 2014; Rui et al., 2015); in addition, mutant HTT has been proposed to compromise selective autophagy (Croce and Yamamoto, 2019). These findings suggest that HTT may be a critical player at different stages of autophagy, such as substrate recognition and trafficking.

#### Supplemental Figures:

**Supplemental Figure 1 (Related to Figure 1): TBK1 phosphorylates HTT specifically at S13 and S16:** (**A**) Mass spectra of recombinant HTT 2-140 23Q after the indicated times of phosphorylation by TBK1. The lower panel is a representative western blot using pT3, pS13 and pS16 specific antibodies of HTT 1-140 23Q after its phosphorylation with TBK1 according to the indicated reaction time. (**B**) Mass spectra of recombinant HTT 1-171 23Q after the indicated times of phosphorylation by TBK1. The lower panel is a representative western blot using pT3, pS13 and pS16 specific antibodies of HTT 1-171 23Q after its phosphorylation with TBK1 according to the indicated reaction time. **(C)** Representative mass spectra of HTTex1 23Q after 16 hours of phosphorylation by TBK1. (**D**) Representative mass spectra of HTTex1 23Q pT3 after 16 hours of phosphorylation by TBK1. (**E**) Representative mass spectra of HTTex1-23Q S13D/S16D after 16 hours of phosphorylation by TBK1 (* salt adducts, + Sinapinic acid matrix adduct).

**Supplemental Figure 2 (Related to Figure 1): TBK1 phosphorylates HTT specifically at S13 S16 much efficiently than IKK β:** (**A**) Representative western blot of HTTex1 phosphorylation by TBK1 and (**B**) phosphorylation by IKKβ using pS13 and pS16 specific antibodies for the indicated with the phosphorylation reaction monitored overtime. (**C**) Representative MALDI spectra of HTTex1 phosphorylation by TBK1 and (**D**) phosphorylation by IKKβ overtime.

**Supplemental Figure 3 (Related to Figure 2): TBK1 phosphorylates longer fragments and full-length HTT in cellular models:** (**A**) MS/MS spectra of the S13/S16 amino acids containing peptides from HTT immunoprecipitated (IP) from HEK293 cell lysates after coexpression of HTTex1 16Q eGFP with TBK1 KD (upper panel) or TBK1 (lower panel). (**B**) Spectral counting of the total WT peptide compared to S13 and S16 peptides from the coexpression with TBK1 or TBK1 KD. **(C)** Representative western blot of the indicated kinase from the immunoprecipitation of HTTex1 16Q eGFP co-expressed with kinase for 48 hours in HEK293 cells. **(D)** Representative western blot of the indicated kinase from the immunoprecipitation of HTTex1 72Q eGFP co-expressed with kinase for 48 hours in HEK293 cells. **(E)** Representative western blot of endogenous HTT and pS13 after treatment with TBK1 siRNA for 72 hours in rat primary striatal neuronal cells. (**F**) Quantification of the fold change in HTT normalized to the HTT levels in non-targeting siRNA normalized to tubulin from the experiments like in E. (**G**) Representative western blot of endogenous HTT and pS13 after treatment with the indicated TBK1 inhibitors (concentrations of 2, 1. And 0.5 µM) for 48 hours in rat primary striatal neuronal cells. (**H**) Quantification of the fold change in HTT compared to DMSO treatment normalized to tubulin from the experiments like in G. (**I**) Western blot of HTT and HTT pS13 levels in cortex tissue lysates from *Tbk1 -/-* and *Tnf-α -/-* mice. **(J)** Quantification of the fold change in HTT normalized to tubulin from I.

**Supplemental Figure 4 (Related to Figure 3): TBK1 overexpression phenocopies effects of HTT S13/S16 phosphomimetic mutations on subcellular localization, aggregation and cytotoxicity of HTT:** (**A**) Immunofluorescence images of the co-expression of HTTex1 16Q eGFP or 72Q eGFP and the indicated kinases for 48 hours in HEK293 cells (scale bar 10µM). (**B**) Quantification of the fold change in nuclear localization upon co-expression of the indicated kinase from experiments like in A. **(C)** Representative western blot of subcellular fractions co-expressing HTTex1 16Q eGFP or 72Q eGFP with the indicated kinases for 16 hours in HEK293 cells. (**D**) Quantification fold change of the ratio of nuclear HTTex1 to cytosolic HTTex1 upon co-expression of the indicated kinase compared to HTTex116Q eGFP from the experiments like in C normalized to respective nuclear and cytosolic Lamin-B and tubulin. (**E**) Representative western blot of soluble HTT, aggregates, pS13 and pT3 upon co-expression of HTTex1 72Q eGFP with the TBK1 or TBK1 KD for 48 hours in HEK293 cells. **(F)** Quantification of soluble and HTT aggregates indicating the fold change compared to TBK1 KD expressing HTTex1 from the experiments like in E. (**G**) Representative immunoblot of HTT and pS13 from a filter retardation assay of HTTex1 43Q fibrils phosphorylated for the indicated times with TBK1 and IKKβ. (**H**) Representative western blot of soluble HTT and HTT pS13 from the zQ175 HD mouse model whole-brain lysate at 24 weeks of age. (**I**) Representative immunoblot of insoluble HTT and pS13 from a filter retardation assay of whole-brain lysate from the zQ175 HD mouse model at 24 weeks of age.

**Supplemental Figure 5 (Related to Figure 5): Downregulation of the *TBK1* worm orthologues ikke-1 and unc-51 using RNAi:** Quantification of the relative mRNA quantity of the indicated worm kinases upon RNAi treatment normalized to act-1.

**Supplemental Figure 6 (Related to Figure 6 and 7): TBK1 overexpression regulates autophagy:** (**A)** Representative immunoblot of filter retardation assay of HTT from the insoluble cellular fraction, HEK293 cells were co-expressing HTTex1 72Q eGFP and TBK1 or TBK1 KD for 48 hours and for last 16 hour before they were treated with indicated proteasome inhibitor (MG132- 5uM) or autophagy inhibitors (Baf-A-200nM, $NH_4Cl$-10mM, 3-MA- 5mM). Quantification of the fold change of HTT aggregates compared to TBK1 KD treated with DMSO normalized to GAPDH from the blots like in bottom. (**B**) Representative immunoblot of p62 from soluble HEK293 fraction over expressing TBK1 or TBK1 KD for 24 hours, cells were treated for last 1 or 4 hour before cell lysis with Baf A1 (500nM) as indicated. (**C**) Quantification of the fold change p62 compared to untreated TBK1 KD at time 0 normalized to GAPDH from the experiments like in B. (**D**) Representative immunoblot of p62 from soluble rat primary striatal cellular fraction overexpressing TBK1 or TBK1 KD for 5 days, cells were treated for last 1 or 4 hour before cell lysis with Baf A1 (500nM) as indicated. (**E**) Quantification graph of the fold change p62 compared to untreated TBK1 KD at time 0 normalized to GAPDH from the experiments like in D. Representative immunofluorescence image showing the colocalization of TBK1 with LC3 (**F**), ubiquitin (**G**) and HTT (H) and p62 (**I**) in rat primary striatal cells at DIV14 (scale bar 5µM). (**J**) Representative western of phospho-OPTN, phosho-p63 and LC3 from HEK293 cells co-expressing HTTex1 72Q eGFP and TBK1 KD, TBK1 or TBK1 Δ 690-713 for 48 hours.

**Supplemental Figure 7 (Related to discussion): TBK1 inhibition affects HTT levels: (A)** Representative immunofluorescence image showing the localization of TBK1 with HTTex1 72Q eGFP in HEK293 cells after 48 hours of co-transfection (scale bar 10µM). **(B)** Representative immunofluorescence image showing the localization of TBK1 with HTTex1 72Q eGFP in rat primary striatal cells at DIV14 after 5 days of co-transfection. (**C**) Quantification of TBK1-positive aggregates in HEK293 cells and rat primary striatal cells from immunofluorescence images similar to those in A and B. (**D**) Quantification of the fold changes in TBK1 and pS172 TBK1 from control and R6/2 brain lysates at 4,8 and 12 weeks of age compared to Wt 4-week control normalized to HSP90. (**E**) Representative western blot showing TBK1 or pTBK1 from postmortem cortical tissue from normal controls and HD patients. (**F**) Quantification of the fold changes in TBK1 and pS172 TBK1 compared to the first normal tissue normalized to actin from E. (**G**) Quantification of the expression (integrated intensity) of

TBK1 pS172 immunoreactivity from middle temporal gyrus tissue microarray containing n=18 control and n=21 HD postmortem tissue samples. (**H**) Representative images of control and HD postmortem tissue showing TBK1 pS172 immunoreactivity from middle temporal gyrus tissue microarray. (**I**) Graph comparing the integrated intensities of TBK1 pS172 and HTT polyQ specific 1C2 immunostaining from middle temporal gyrus tissue microarray of HD and control cases.

#### Materials and methods

Antibodies and chemicals

The antibodies and chemical used in the study were listed in Table1 including the clone name, catalog numbers, providers.

Plasmids and Constructs

cDNAs encoding N-terminal HTT fragments (exon 1) bearing different polyQ lengths (Q16, Q39, or Q72) and HTT N548 were described previously1. Myc-TBK1 plasmid (RC205238) and Myc-IKKβ plasmid (RC219154) were obtained from Origene, Rockville, MD, USA. TBK1 Δ690-713 was obtained by site directed mutagenesis of RC205238 plasmid. Flag-TBK1 kinase dead (K38A) plasmid (DU12696) was obtained from MRC PPU Reagents and Services, School of Life Sciences, University of Dundee. TBK1 lentiviral plasmid (LV330421) was obtained from Abmgood (Applied Biological Materials Inc) Richmond, Canada. TBK1 KD (K38A) lentiviral plasmid was obtained by site directed mutagenesis mutation to LV330421 plasmid. To obtain plasmids of TBK1 and TBK1 kinase dead for C. elegans expression, TBK1 and TBK1 kinase dead sequences were PCR amplified from RC205238 and DU12696 and introduced to pPD30_38 (Addgene Plasmid #1443).

##### *In vitro* phosphorylation of HTTex1 with TBK1 and IKKβ

HTTex1 proteins were expressed, purified, disaggregated and prepared for phosphorylation as we previously described (Ahmad et al., 2016; Reif et al., 2018). After the disaggregation of HTTex1 and evaporation of any residual TFA acid, the protein was dissolved at a final concentration of 0.5 mg/ml, in the phosphorylation buffer: 50 mM Tris, 25 mM $MgCl_2$, 8 mM EGTA, 4 mM EDTA and 1 mM DTT (This solution is to be freshly prepared each time). Next, the pH of the reaction was adjusted to 7.4, and 5 mM Mg-ATP was added to the reaction solution, as well as purified recombinant TBK1 or IKKβ (# DU12469, #DU3167 -MRC PPU Reagents and Services, University of Dundee) at a ratio of 1μg of kinase per 30 μg of protein. The reaction mixture was kept at 30°C and monitored overtime by mass spectroscopy (LC-ESI-MS) and western blot (WB).

To perform the Mass spectroscopy, 5 μL of the reaction mixture was injected in the Thermo Scientific LTQ ion trap. To obtain the intact molar mass, the different charges states were deconvoluted using Magtran software. Additionally, 10 μL of the kinase reaction solution was supplemented with 2X Laemmli loading buffer, snap frozen and stored in 20°C for further Immunoblotting Assay (see below)

**HEK293, HEK293T Cell Culture, transfections**

HEK293 and HEK293T cells were cultured t 95% air and 5% CO2 in Dulbecco's Modified Eagle Medium (Gibco) supplemented with 10% Fetal bovine serum (Gibco), Penicillin-Streptomycin Thermo Fisher. Transfections were carried by Lipofectamine 2000 according to the manufacturers protocol. Lysis was performed in RIPA lysis buffer (150 mM sodium chloride, Triton X-100, 0.5% sodium deoxycholate, 0.1% SDS (sodium dodecyl sulfate) ,50 mM Tris, pH 8.0) or NP40 lysis buffer (150 mM sodium chloride 1.0% NP-40, 0.5% sodium deoxycholate, 0.1% SDS (sodium dodecyl sulfate), 50 mM Tris, pH 8.0 supplemented with 1× protease and phosphatase inhibitor 2, 3 mixture (Sigma). Cell lysates were then centrifuged at 15000xg for 20 min and supernatant was collected as soluble fraction. Pellet was washed twice with lysis buffer and then re-suspended with 500ul of lysis buffer supplemented with 2% SDS. Pellet was sonicated for 9 second with 3 sec on and 3 sec off at 60% amplitude. Protein concentration was measured using BCA system and about 20-60 ug of protein is processed for WB or filter retardation assay.

Primary neuronal cell culture and protein expression and siRNA treatment inhibitor treatment

Primary cultures of medium spiny neurons were prepared under the local experimental license (VD2137 and VD3392). Briefly, P0 rat pups were dissected and striatal cells were isolated and dissociated by repeated pipetting. Cells were pelleted by centrifugation, resuspended in the culture medium [Neurobasal medium (Invitrogen) complemented with 1% B27 (Invitrogen), Pen-Strep, L-Glutamine and KCl], plated in poly-L- lysine-coated multi-well culture dishes at a density of 150 000 cells/ cm2 and cultured at 37°C in 5% CO2/air atmosphere. Half the medium was replaced on DIV 4, and half of the medium was replaced weekly thereafter. For lentiviral- mediated protein expression, cultures were infected on day *in-vitro* (DIV) 7. Plasmid transfections were carried by Lipofectamine 2000 according to the manufacturers protocol on. The siRNA transfections were carried by Lipofectamine 2000 according to the manufacturers protocol.

**Immunoblotting Assay and Filter Retardation Assay**

Cells were washed twice with PBS and harvested in either RIPA buffer (20 mM Tris-HCl (pH 8), 150 mM NaCl, 1 mM Na2 EDTA, 1 mM EGTA, 1% Triton X-100, 1% sodium deoxycholate- with protease (Sigma) and phosphatase inhibitors (cocktail 2 and 3 Sigma) or NP40 lysis buffer (20 mM Tris-HCl (pH 8), 150 mM NaCl, 1 mM EDTA, 1 mM EGTA, 1% NP40, 1% sodium deoxycholate- with protease (Sigma) and phosphatase inhibitors (cocktail 2 and 3 Sigma). Lysates were incubated on ice for 30 min and vortexed every 10 min. Protein extracts were cleared of cell debris by centrifugation at 20000 g for 15 min at 4°C. Samples were divided in two and were either collected at insoluble pellet fraction, or spun homogenates. Insoluble pellet fractions were washed with RIPA buffer and sonicated in ul RIPA supplemented 2% SDS with settings 40% amplitude for 9s (3s on; 3s off) and collected as detergent insoluble fraction and used in filter retardation analysis of HTT aggregates. The protein concentration of the fractions was determined using BCA kit (Thermo Fisher). 30 ug of total or spun protein in Laemmli loading buffer was denatured at 85 °C for 5 min and separated by sodium dodecyl sulfate–polyacrylamide gel electrophoresis (SDS–PAGE) on a 7 to 15% polyacrylamide gel. Proteins were subjected to WB using nitrocellulose membrane. After blocking using Odyssey Blocking Buffer (PBS) (P/N: 927-40000) Li-Cor, Lincoln, Nebraska USA, primary antibodies (Table 1) incubation was carried out overnight at 4 °C, and secondary antibody incubations for 1 h at room temperature. Protein bands were detected using Odyssey® CLx Imaging System, Li-Cor, Lincoln, Nebraska USA. Filter-retardation assay of insoluble fractions were carried out as described previously(Ochaba et al., 2018; Sontag et al., 2012). 30 µg proteins were passed through a 96-well vacuum filtration apparatus (#1706545; Bio-Rad Laboratories) containing a 0.2-µM cellulose acetate membrane filter (catalog #10404180; Whatman) prewetted in PBS/2% SDS. After three washing steps with PBS/2% SDS, the membrane was fixed, blocked, and incubated with antibodies as previously described. Immunodetected proteins quantified using the ImageJ software.

**Immunoprecipitation**

Immunoprecipitation was performed using Dynabeads Protein G (catalog #10004D; Thermo Fisher Scientific) following the manufacturer's instructions and using an HTT-specific antibody (D7F7) or ab105119. The pulled-down material was analyzed by SDS/PAGE and WB.

**Immunocytochemistry**

For immunostaining, cells were grown on poly-L-lysine- coated glass coverslips, infected as described above with different expression vectors. Three days post infection; cells were fixed with 4% paraformaldehyde in PBS (pH 7.4) for 15 min. After fixing, cells were washed three times with ice-cold PBS and blocked with 10% NGS (Invitrogen) in PBS supplemented with

0.1% Triton X- 100 (Sigma). Primary antibody diluted 1:100 in PBS with 5% NGS and 0.1% Triton X-100, and incubated with the cells overnight at 4°C. Cells were washed three times with PBS and incubated at RT with secondary donkey anti-mouse IgG Alexa 568 (Invitrogen, Carlsbad, CA, USA) in PBS with 1% NGS, followed by three washes with PBS. Images were taken using the same inverted confocal laser scanning microscope with Polyvinyl alcohol mounting medium with DABCO (Sigma-Aldrich, St. Louis, Missouri, USA) immersion 63· objective as above. For thioflavin S staining was carried out on fixed cells by incubating with 0.05% thioflavin S (Sigma) for 8 min followed by wash with 80% ethanol for 5 min (three times) each before the antibody incubations.

**Transfection of primary neurons**

Neurons were co-transfected using Lipofectamine 2000 (Thermo Fisher Scientific) at day *in-vitro* (DIV) 5 or 6. Per well of 24-well plates, 1 ug of HTT plasmid, and 0.1 ug of GFP was used with a ratio DNA/Lipofectamine of 1/1.75. DNA and Lipofectamine were diluted separately in 50ul of OptiMEM/ well for each and incubated for 5 min before being combined. After being combined, the transfecting solution is incubated for 90 min at room temperature. During the incubation of the transfecting solution, the plates are emptied, and the medium is replaced with 400 ul of fresh OptiMEM. After the incubation, 100 ul of transfection solution is deposited in each well. The plates are returned to the incubator for 4.5 h when the wells are empty, and the medium was replaced with fresh Neurobasal medium.

**Cytoplasmic and nuclear fractionation**

The fractionation was carried out as described previously (Have et al., 2012) adapting to the cell density. 48-hour post transfection HEK293 cells from 10cm culture dish were washed 3x times with PBS scraped in PBS (2ml) and cells were pelleted by centrifuging for 4mins at 200g.

Cell pellet was resuspended in 1ml HEPES lysis buffer (HEPES, Ph 7.9, 10mM. MgCl2 1.5mM, KCl 10mM, DTT 0.5mM supplemented with protease inhibitor cocktail and phosphatase inhibitor cocktail 2 and 3). Resuspended cells were then transferred into a pre-chilled 7ml Dounce homogenizer and cells were sheared using 10 strokes of a tight pestle. Centrifuged the Dounced cells for 5 mins at 4ºC, 1000rpm. Retained supernatant as cytoplasmic fraction. Re-suspended nuclear pellet in 1ml of Sucrose 1 (0.25M Sucrose, 10mM MgCl2) Cushioned on top of layer of 1 ml Sucrose 2 (0.35M Sucrose, 0.5mM MgCl2) by slowly pipetting solution Sucrose 1 on top of Sucrose 2. Centrifuged for 10 mins at 4ºC, 3500rpm and retained pellet as nuclear fraction. Nuclear fraction was resuspended 500ul RIPA buffer and sonicated for 9s (3s on 3s off, at 60 % amplitude) and samples were collected as nuclear lysate. The cytoplasmic and nuclear fraction protein cotes were estimated by BCA and the levels of HTT were monitored WB using HTT antibody, β-Tubulin and Lamin-B.

**Nuclear condensation cell death assay**

The toxicity experiments were performed in primary cortical and primary striatal neurons according to our established protocol(Arbez et al., 2017). Nuclei were stained using Hoechst 3342 (Sigma; 0.2 μg/ml in PBS for 5 min). Automated picture acquisition was performed using a Zeiss Axiovert 200 inverted microscope with a 10× objective. Automatic quantification of the nuclear intensity of transfected cells was performed using Volocity. The cells were considered dead when their nuclear intensity was higher than the average intensity plus two standard deviations. Each condition was performed in quadruplicate within each experiment, and each experiment was repeated in at least four independent neuronal preparations for each construct studied.

**TUNEL assay**

DNA fragmentation was detected using the TUNEL method as described previously (Mahul-Mellier et al., 2015). Primary neurons were co-transfected for with HTTex1 16Q or 72Q eGFP along with TBK1 or TBK1 KD for indicated time. Then, the cells were washed 3 times with PBS to remove unattached recombinant α-syn and fixed in 4% PFA for 15 min at 4 °C. Cells were permeabilized in a solution composed of 0.1% Triton X-100 in 0.1% citrate buffer, pH 6.0, and then washed in PBS buffer before incubation with terminal deoxynucleotide transferase (In Situ Cell Death Detection kit; Roche) for 1 h at 37 °C in a solution containing TMR red dUTP. The neuronal nuclear marker NeuN and TBK1 was stained and the cells were washed and stained with secondary antibodies. Then washed 3 times in PBS and mounted using polyvinyl alcohol (PVA) mounting medium (Sigma-Aldrich). The HTT and kinase co-transfected TUNEL positive neurons were counted and plotted.

**Singulex Assay**

A total of 50 µL/well of dilution buffer (6% BSA, 0.8% Triton X-100, 750 mM NaCl, and complete protease inhibitor) was added to a 96-well plate (catalog #P-96–450V-C; Axygen). Samples to be tested were diluted in artificial cerebral spinal fluid (0.3 M NaCl; 6 mM KCl; 2.8 mM CaCl2-2H2O; 1.6 mM MgCl2-6H20; 1.6 mM Na2HPO4-7H2O; 0.4 mM NaH2PO4-H2O) supplemented with 1% Tween-20 and complete protease inhibitor in a final volume of 150 µL/ well. Finally, 100 µL/well of the MW1 antibody coupled with magnetic particles (appropriately diluted in Erenna Assay buffer, catalog #02–0474-00; Singulex) was added to the assay plate and incubated for 1 h at room temperature under orbital shaking. The beads were then washed with Erenna System buffer (catalog #02–0111-00; Singulex) and resuspended using 20 µL/well of the specific detection antibody labeled with D2 fluorophore (or Alexa-647 fluorophore) appropriately diluted in Erenna Assay buffer. The plate was incubated for 1 h at room temperature under shaking. After washing, the beads were resuspended and transferred to a new 96-well plate. A total of 10 μL/well of Erenna buffer B (catalog #02–0297-00; Singulex) was added to the beads for elution and incubated for 5 min at room temperature under orbital shaking. The eluted complex was magnetically separated from the beads and transferred to a 384- well plate (Nunc catalog #264573; Sigma-Aldrich) where it was neutralized with 10 μL/well of Erenna buffer D (catalog #02–0368-00; Singulex). Finally, the 384- well plate was heat-sealed and analyzed with the Erenna Immunoassay System.

**MS/MS analysis**

HTT immunoprecipitated was loaded on to SDS-PAGE the bands corresponding to HTT were cut and In-gel digestion was carried out. Briefly gel bands were washed 2x with 100 ul 50% ACN/100 mM NH4CO3; 1x 50 ul CAN, 100 ul of 10 mM DTT in 50 mM TEAB was added and incubated for 45 min, 37°C, removed DTT, added 100 ul of 15 mM IAM solution in 50 mM TEAB, incubated for 1h, RT, in darkness, removed IAM added 20 ul Chymotrypsin (50 ng/ul), incubated 5 min, added 40 ul TEAB buffer to digest overnight. Digested peptides were extracted by 150ul 0.1% TFA / 50% ACN Samples were dried and dissolved in 20uL of 2.5% ACN+0.1% FA. Then 3 μL sample was injected into nanoUPLC (Waters) and MS measurement were performed by QExactive (Thermo). All raw data were search against Swissprot via Mascot and search against database containing HTTex1 sequence via Byonic 3.5 with the consideration of oxidation of methionine, phosphorylation of serine/ threonine, and acetylation of protein N-terminal as variable modifications.

**Lentiviral vector production**

SIN-W-PGK and SIN-W-tetracycline responsive element (TRE) expression vectors were used for the production of lentiviruses in human cells (HEK293T) using a four-plasmid system as described previously (1). Viral pellets were resuspended in phosphate-buffered saline (PBS) with 0.5% bovine serum albumin and stored at 80°C. Prior to the utilization, viral stocks were diluted with cell culture medium to a concentration of 1500 ng p24 antigen/mL as measured by ELISA (RETROtek, Gentaur, Kampenhout, Belgium), and applied to the cell cultures at the concentration of 50 ng p24/mL of culture medium for each virus.

**_C. elegans_ RNAi treatment and biochemical analysis**

Bacterial clones for RNAi (cst-1, cst-2, unc-51 and ikke-1) were obtained from the Ahringer RNAi library (HTTps://www.sourcebioscience.com/products/life-science-research/clones/rnai-resources/c-elegans-rnai-collection-ahringer/). RNAi clones were grown on LB with ampicillin (100 ug/mL) and tetracycline (12.5ug/mL) o/n. Then grown up from a 1/10 dilution in LB with ampicillin (100ug/mL) for 7h which were used to spot medium treatment plates 6 drops each plate, evenly dispersed across the plate. Worm strains are maintained on standard NGM agar plates for multiple generations growing on OP50. Medium NGM treatment agar plates were prepared for all RNAi clones for both Q15 and Q128 worms in duplicate. Post 24h incubation of the RNAi clones on the NGM treatment plates 10 L4's of the corresponding strain was placed on RNAi treatment plates. Post-maturation and egg laying overnight the L4's were removed ~18h later. Worms were harvested ~72h after removal of matured worms & hatching of the laid eggs. Worms washed off the plate using M9 and centrifuged at a 2000rpm for 1 min to form a live worm pellet. A small portion of worms were used for isolation of total mRNA using the Trizol. The Isolated mRNA were transcribed and q-PCR was carried out to determine the relative mRNA level using primers of act-1, ikke-1 and unc-51. Other portion of worms were flash frozen in liquid nitrogen. 200 ul RIPA buffer (20 mM Tris-HCl (pH 8), 150 mM NaCl, 1 mM Na2 EDTA, 1 mM EGTA, 1% Triton X-100, 1% sodium deoxycholate- with protease (Sigma) and phosphatase inhibitors (cocktail 2 and 3

Sigma) was added to the frozen pellet and allow the pellet to thaw on regular ice just until melted. Then the contents of the tube were sonicated with microtip using the settings; 40% amplitude for 30 s (15s on; 45s off) (Zanin et al., 2011), and then centrifuges at 15000 rpm for 20 mins and the supernatant was collected as soluble protein and used for analysis using the SDS-PAGE followed by WB. The pellets were washed with RIPA buffer and sonicated in 200 ul RIPA supplemented 2% SDS with settings 40% amplitude for 30 s (15s on; 45s off) and collected as detergent insoluble fraction and used in filter retardation analysis of HTT aggregates.

**Transgenic TBK1 KD and TBK1 *C. elegans* lines**

C. elegans were cultured according the standard methods (Lee et al., 2017). In short, animals were maintained at 20°C or 15°C on NGM agar plates seeded with E. coli (OP50). The Punc-54HTT513(Q15)::YFP or Punc-54HTT513(Q128)::YFP were previously published and expresses HTT N513 tagged with YFP transgene in body wall muscle cells (Lee et al., 2017). To generate transgenic TBK1 and TBK1 KD animals, 50ng/µL of DNA encoding Punc-54 TBK1 or Punc-54 TBK1 KD were microinjected along with Punc-54 DS-Red selection marker into the gonads of adult wild type hermaphrodites to generate multiple (at least 3) independent lines transmitting extrachromosomal arrays.

***C. elegans* Motility assay**

Individual lines co-expressing HTT N513 15Q or 128Q with TBK1 or TBK1 KD were examined at days 5 of adulthood. A single L4 worm of each line was grown in NGM agar plate with OP50 bacteria following standard procedures. Three days after pretreatment 30 L4 worms were picked from each line and placed on fresh NGM agar plates containing 5-FU to avoid progeny development. At $5^{th}$ day of adulthood the video was recorded for each individual line and the motility was measured in speed (mm/s) using the wrMTrck plugin using the ImageJ as described previously (Au - Nussbaum-Krammer et al., 2015).

##### *C. elegans* Lifespan assay

Animals were acclimated at 20°C for at least two generations before initiating lifespan analyses. Forty age-matched L4 larvae were placed on NGM plates seeded with E. coli (OP50) and cultured at 20°C. During the reproductive time of their life cycle, animals were transferred away from their progeny by daily passaging to freshly seeded plates. Post-reproductive adults were passaged as needed to prevent starvation. Animals were scored as deceased when they no longer moved upon prodding the head several times with a platinum wire as described previously (Lee et al., 2017). Animals that crawled off the plate were censored in statistical analyses by adjusting the total population to the number of animals seen on the plate on a given day.

##### Detection of TBK1 from R6/2 HD mouse brain tissue

4-, 8-, or 12-wk n=3 / genotype brain hemispheres; Wild type or R6/2 transgenic half brains were homogenized by mechanical shearing (Polytron PT 1200 E) for 1X 30 sec in 1 ml KCL buffer (50 mM Tris–HCl pH 8.0, 150 mM KCl, 5 mM EDTA, 10% Glycerol with 10 mM dithiothreitol, 1 mM phenylmethylsulfonyl fluoride, ‘complete’ protease inhibitors (ROCHE), and PhosSTOP (ROCHE), at 4 °C. Samples were divided in two and were either collected at TOTAL homogenates, or further spun 2X at 13,000 X g for 20 min at 4 °C as SPUN homogenates. The protein concentration of the total and spun (supernatant) fractions were determined using BCA kit (Thermo Fisher). 30 ug of total or spun protein in Laemmli loading buffer was denatured at 85 °C for 5 min and separated by 10% Criterion TGX Stain-free SDS-PAGE (Bio-Rad). Membranes were blocked for 1 hr at RT in 5% non-fat dried milk in PBST

(TBS, 0.2% Tween-20), and then incubated with gentle agitation overnight at 4 °C with the primary antibody (in TBST with 1% non-fat dried milk). Blots were immunoprobed with either: TBK1 (Cell Signaling Technologies #3504=1:1000) and TBK1 pS172 (Cell Signaling Technologies #5483=1:1000) For chemilluminescent detection, blots were washed 3X in TBST for 5 min, probed with HRP-linked secondary antibodies (in TBST with 0.5% non-fat dried milk) for 1 hr at RT and washed 3X in TBST. Protein was detected by chemilluminescense (Clarity, Bio-Rad) according to the manufacturer's instructions. The signals were captured on a ChemiDoc Touch Imaging System (Bio-Rad), then subsequently quantified using Image Lab v5.2.1 (Bio-Rad).

**Detection of TBK1 from Human samples mouse brain tissue**

The post-mortem human brain tissue used in this study was obtained from the Cambridge Brain Bank, Division of the Human Research Tissue Bank, Addenbrooke's Hospital, Cambridge UK. All protocols in this study were approved by Cambridge Brain Bank NRES approval 10/H0308/56. The anonymized human brain cortical tissue 100 mg (n=5 of HD and Control) tissue were homogenized by mechanical shearing using the 30 strokes in Dounce homogenizer in 1 ml KCL buffer (50 mM Tris–HCl pH 8.0, 150 mM KCl, 5 mM EDTA, 10% Glycerol) with 10 mM dithiothreitol, 1 mM phenylmethylsulfonyl fluoride, 'complete' protease inhibitors (Sigma), and Phosphatase inhibitor cocktail 2 and 3 (Sigma). Samples were collected after a spin at 13,000 X g for 20 min at 4 °C. The protein concentration of the total and spun (supernatant) fractions were determined using BCA kit (Thermo Fisher). 50 ug of protein in Laemmli loading buffer was denatured at 85 °C for 5 min and separated by 10% Tris-Glycine SDS-PAGE (Bio-Rad). TBK1 and TBK1pS172 were immunodetected using the Odyssey Infrared Imaging System (LI-COR) and quantified using the ImageJ software.

**Human brain tissue microarrays**

The post-mortem human brain tissue used to construct tissue microarrays (TMAs) in this study was obtained from the Neurological Foundation of New Zealand Human Brain Bank in the Centre for Brain Research, University of Auckland. All protocols in this study were approved by the University of Auckland Human Participants Ethics Committee (2008/279 and 011654), and all families provided informed consent. Human brain TMAs were constructed from paraffin-embedded middle temporal gyrus blocks, using methods described previously (Coppieters et al., 2014; Narayan et al., 2015; Singh-Bains et al., 2019). The TMA comprised a total of 56 2-mm-diameter core samples, 28 from neurologically normal control cases and 28 from HD cases. Final sample sizes used for immunohistochemistry with TBK1 pS172 was n=21 HD and n=18 control cases, which reflects the loss of samples from the slide during processing.

**Immunohistochemistry to detect TBK1 pS172 and 1C2 immunoreactivity in human brain tissue microarrays**

TMA sections were cut at a thickness of 7µm and were annealed to slides by heating at 60°C for 1 hour. Sections were subsequently dewaxed using xylene immersion in xylene twice (1 hour and 10 min, respectively) and rehydrated using a standard graded ethanol series procedure. For immunohistochemistry, slides were immersed in Tris-EDTA retrieval buffer (pH9) (for TBK1 pS172) or sodium citrate buffer (pH6) (for 1C2), heated at 121°C for 2 hours in 2100 Antigen Retriever (Aptum, Pick Cell Laboratories), and then rinsed three times for 5 min in milliQ$H_2O$. For 1C2 immunohistochemistry, once the slides had cooled post-antigen retrieval, an additional incubation in 99% formic acid was carried out for 5 minutes followed by 3 washes in milliQ $H_2O$. Slides were subsequently incubated with an endogenous peroxidase blocking solution (50% methanol, 1% $H_2O_2$, diluted in mQ$H_2O$) for 20 min at room temperature. This was followed by three washes in PBS, and then slides were incubated with blocking buffer (10% normal goat serum in PBS) for 1 hour at room temperature. Primary antibodies rabbit Phospho-TBK1/NAK, Ser172 (1:100, Cell signalling) and mouse 1C2 (1:5000, CAT # MAB1572, Millipore) were applied for incubation overnight at 4°C.

To detect antibody binding, slides were washed in PBS with Triton X-100 for 5 min and then twice in PBS for 5 min each and incubated with biotinylated goat anti-mouse or anti-rabbit antibodies (Sigma-Aldrich) for 3 hours at room temperature. After further washing, they were incubated for 1 hour at room temperature with Sigma Extravidin peroxidase at 1:1000 dilution and then washed and incubated with the peroxidase substrate [3,3′-diaminobenzidine with 0.04% $Ni(NH_4)_2(SO_4)_2$] to develop the colour change. Following PBS-milliQ$H_2O$ washes (3 × 5 min each), the slides were dehydrated in a graded ethanol series followed by xylene and mounted under coverslips with DPX mounting medium (Cat No 06522, Sigma-Aldrich).

Images were acquired for each immunolabelled TMA slide using a VSlide automated slide scanning microscope (Metasystems) running Metafer4 software (version 3.12.133) utilising a TMA imaging protocol outlined by Singh-Bains et al and Narayan et al (Narayan et al., 2015; Singh-Bains et al., 2019). The images acquired were analysed using Metamorph software (Metamorph Offline v.7.8.0, Molecular Devices). Image analysis for the TMA cores were conducted using the ‘Count Nuclei algorithm’ (Narayan and Dragunow, 2010). For each immunolabel, Count Nuclei measurements logged within an excel spreadsheet included the expression of the immunolabel, represented by the integrated intensity. Data obtained from the Metamorph software were statistically analysed using GraphPad Prism (version 7.03). The comparisons between HD and control groups for TMA data were conducted using a 2-tailed Student’s unpaired t-test with Welch’s correction for unequal variances. The correlations between 1C2 and TBK1 pS172 were conducted with Pearson’s correlation analysis (2-tailed).

**Statistics**

The bar graphs plotted represent mean and SEM. Students t-test was used to indicate the significance as < 0.001 - ***, 0.001 to 0.01 **, 0.01 to 0.05 *, ≥ 0.05 ns.

Table 1:

List of antibodies and reagents used in the study.

| **Antibody/reagents** | **Cat No** | **Provider** |
|---|---|---|
| Huntingtin (D7F7) XP® Rabbit mAb | 5656 | Cellsignlaing |
| Recombinant Anti-Huntingtin antibody [EPR5526] (ab109115) | ab109115 | Abcam |
| Anti-Huntingtin Antibody, a.a. 115-129 MAB5490 | MAB5490 | Millipore |
| Anti-Huntingtin Protein Antibody, a.a. 181-810 MAB2166 | MAB2166 | Millipore |
| Anti-Huntingtin Antibody, a.a. 1-82 MAB5492 | MAB5492 | Millipore |
| TBK1/NAK (D1B4) Rabbit mAb #3504 | 3504 | Cell signlaing |
| Phospho-TBK1/NAK (Ser172) (D52C2) XP® Rabbit mAb #5483 | 5483 | Cell signlaing |
| c-Myc Antibody (9E10) | sc-40 | Santa Cruz |
| Anti-NeuN Antibody, clone A60 | MAB377 | Millipore |
| Ubiquitin P4D1 | Sc-8017 | Santa Cruz |
| GAPDH /14C10) | 2118S | Cell signaling |
| Huntingtin EM48 | MAB5374 | millipore |
| Optineurin (C-Term) Polyclonal Antibody | № 100000 | Cayman Chemical |
| Anti-LC3B antibody (ab48394) | ab48394 | Abcam |
| pT3-Huntingtin | CHDI-90001528-2 | CHDI/Thermo Scientific |
| SQSTM1 monoclonal antibody (M01), clone 2C11 | H00008878-M01 | Abnova |
| Anti-p62 (SQSTM1) S403 | D343-3 | MBL |
| pS13 Huntingtin | CHDI-90001039-1 | CHDI/ Thermo Scientific |
| Phospho-Optineurin (Ser177) Antibody | 57548 | Cellsignaling |
| Anti-beta tubulin | ab6046 | Abcam |
| pS16-Huntingtin | ZCH11020 | Lashuel Lab |
| Amlexanox | 4857 | Tocris |
| BX 795 | 4318 | Tocris |
| MRT 67307 dihydrochloride | 5134 | Tocris |
| MRT 68601 hydrochloride | 5067 | Tocris |
| **siRNAs** | | |
| Name | Sequence | |
| Nontar 1 | AUGAACGUGAAUUGCUCAA | Microsynth |
| Nontar 2 | UAAGGCUAUGAAGAAUAC | Microsynth |
| Nontar 3 | AUGUAUUGGCCUGUAUUAG | Microsynth |
| Rat siTBK1-1 | GGGAACAUCAUGCGCGUCA | Microsynth |
| Rat siTBK1-2 | CUAGAGAGUUAGAGGACGA | Microsynth |

Sup. Fig. 1

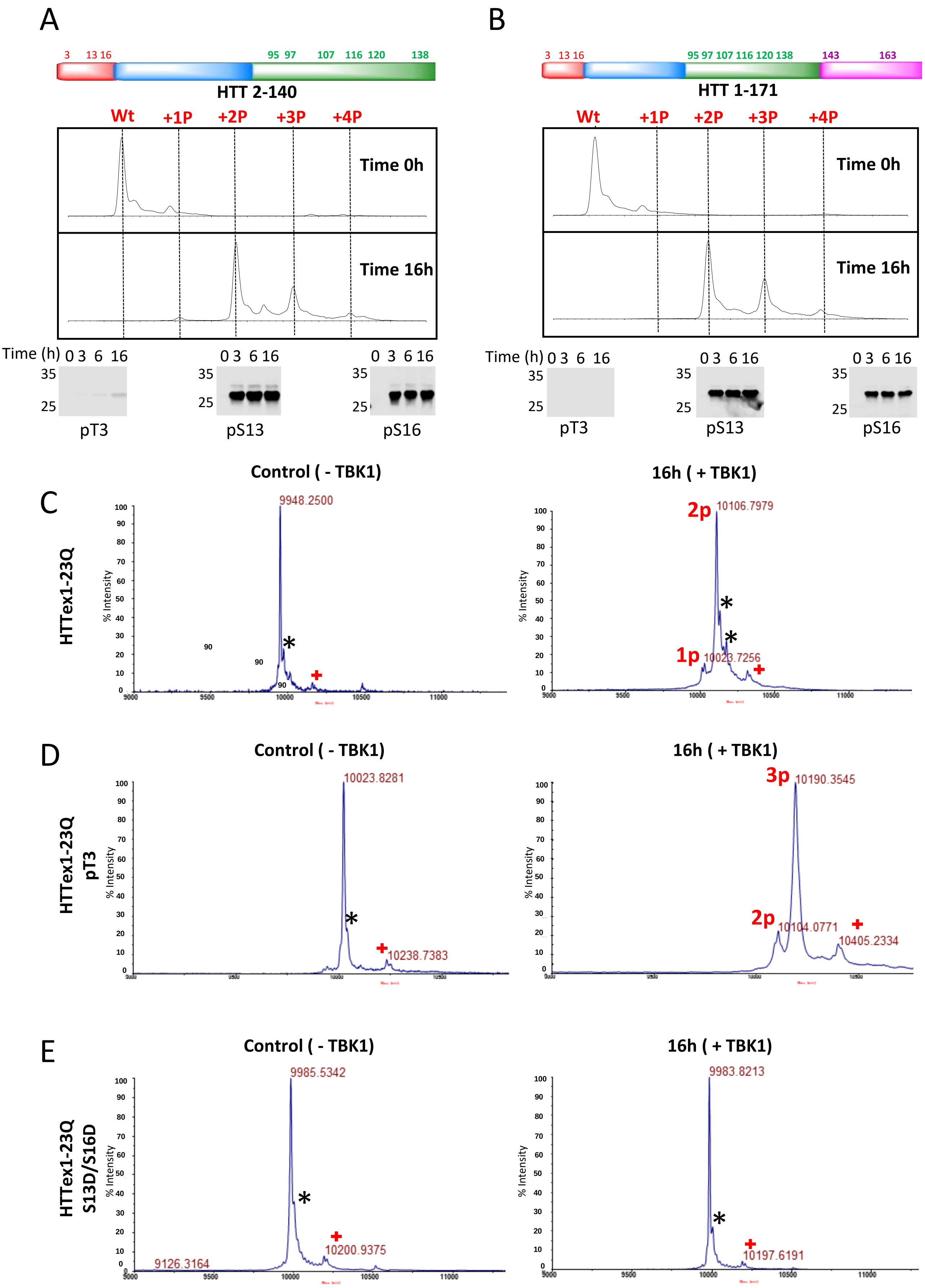

A
3 13 16 95 97 107 116 120 138
HTT 2-140
Wt +1P +2P +3P +4P
Time 0h
Time 16h
Time (h) 0 3 6 16
35
25
pT3
pS13
pS16
B
3 13 16 95 97 107 116 120 138 143 163
HTT 1-171
Wt +1P +2P +3P +4P
Time 0h
Time 16h
pT3
pS13
pS16
C
HTTex1-23Q
Control ( - TBK1)
16h ( + TBK1)
9948.2500
2p 10106.7979
1p 10023.7256
% Intensity
D
HTTex1-23Q pT3
Control ( - TBK1)
16h ( + TBK1)
10023.8281
10238.7383
3p 10190.3545
2p 10104.0771
10405.2334
E
HTTex1-23Q S13D/S16D
Control ( - TBK1)
16h ( + TBK1)
9985.5342
10200.9375
9126.3164
9983.8213
10197.6191

Sup. Fig. 2

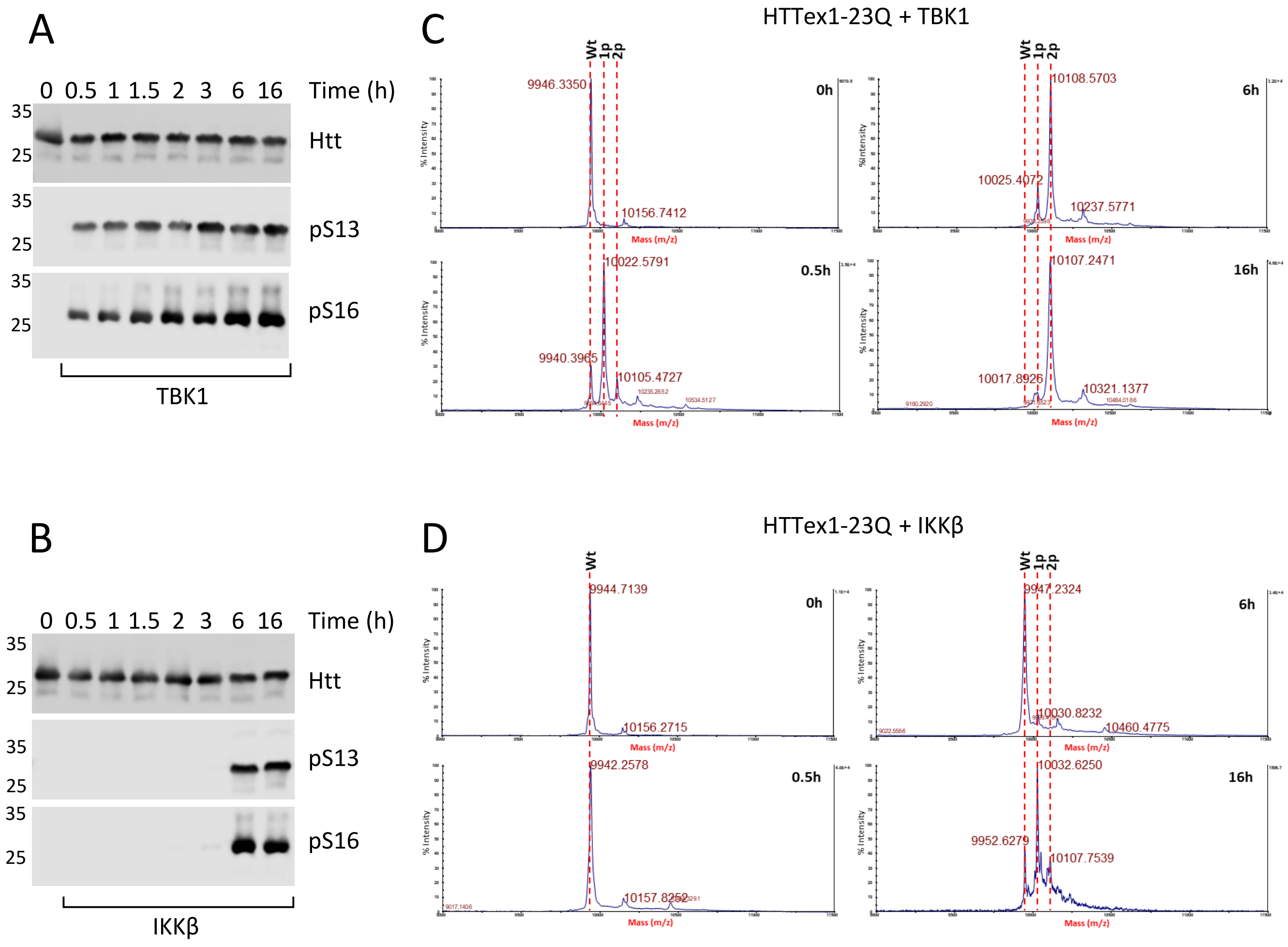
A
0 0.5 1 1.5 2 3 6 16 Time (h)
35
25
Htt
pS13
pS16
TBK1
B
0 0.5 1 1.5 2 3 6 16 Time (h)
Htt
pS13
pS16
IKKβ
C
HTTex1-23Q + TBK1
Wt
1p
2p
9946.3350
10156.7412
0h
10108.5703
10025.4072
10237.5771
6h
10022.5791
9940.3965
10105.4727
0.5h
10107.2471
10017.8926
10321.1377
16h
Mass (m/z)
% Intensity
D
HTTex1-23Q + IKKβ
9944.7139
10156.2715
0h
9947.2324
10030.8232
10460.4775
6h
9942.2578
10157.8252
0.5h
10032.6250
9952.6279
10107.7539
16h

Sup. Fig. 3

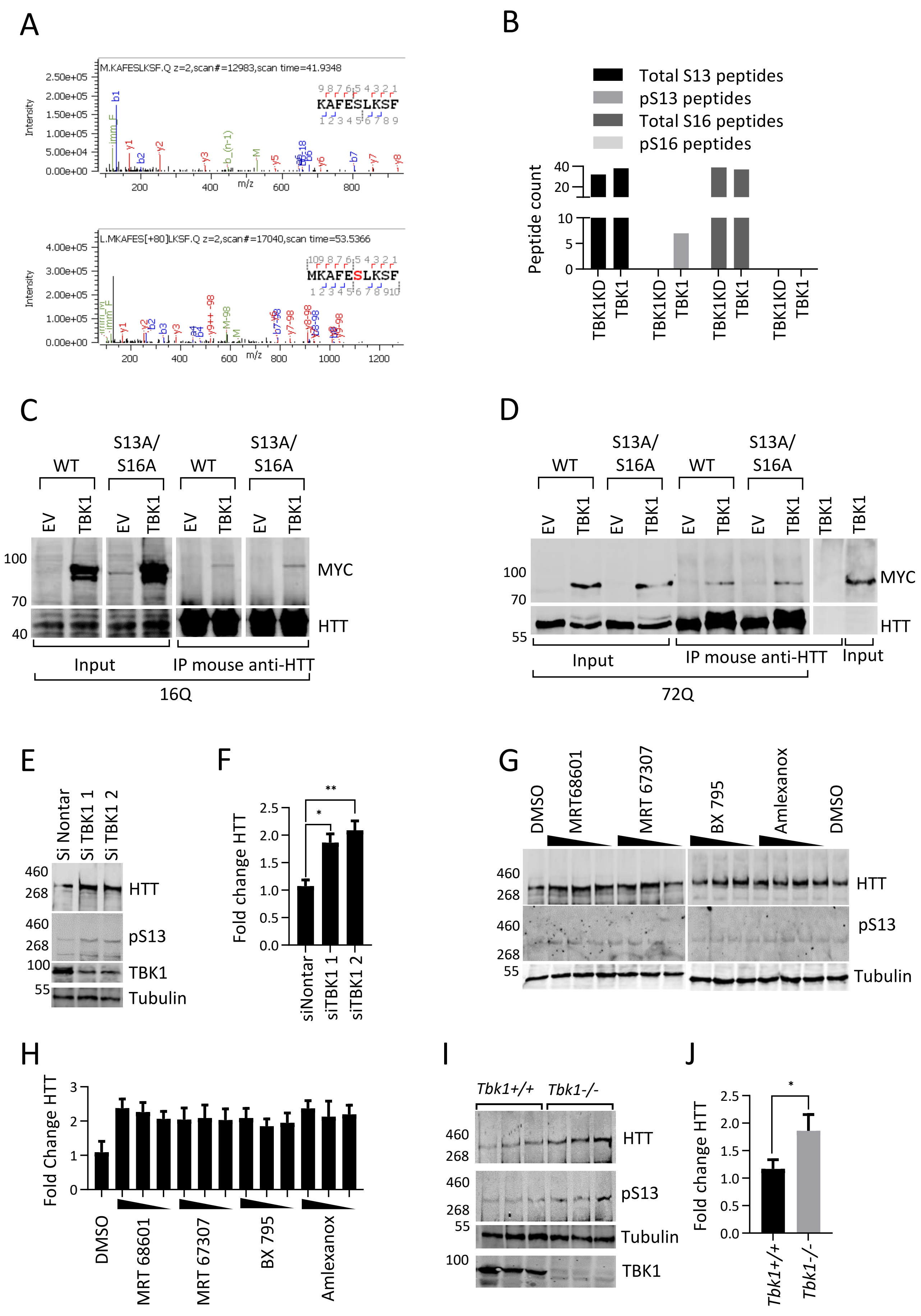

A
M.KAFESLKSF.Q z=2,scan#=12983,scan time=41.9348
KAFESLKSF
Intensity
m/z
L.MKAFES[+80]LKSF.Q z=2,scan#=17040,scan time=53.5366
MKAFESLKSF
B
Total S13 peptides
pS13 peptides
Total S16 peptides
pS16 peptides
Peptide count
TBK1KD
TBK1
C
WT
S13A/ S16A
EV
TBK1
MYC
HTT
Input
IP mouse anti-HTT
16Q
D
WT
S13A/ S16A
EV
TBK1
MYC
HTT
Input
IP mouse anti-HTT
Input
72Q
E
Si Nontar
Si TBK1 1
Si TBK1 2
HTT
pS13
TBK1
Tubulin
F
Fold change HTT
siNontar
siTBK1 1
siTBK1 2
G
DMSO
MRT68601
MRT 67307
BX 795
Amlexanox
DMSO
HTT
pS13
Tubulin
H
Fold Change HTT
DMSO
MRT 68601
MRT 67307
BX 795
Amlexanox
I
Tbk1+/+
Tbk1-/-
HTT
pS13
Tubulin
TBK1
J
Fold change HTT
Tbk1+/+
Tbk1-/-

Sup. Fig. 4

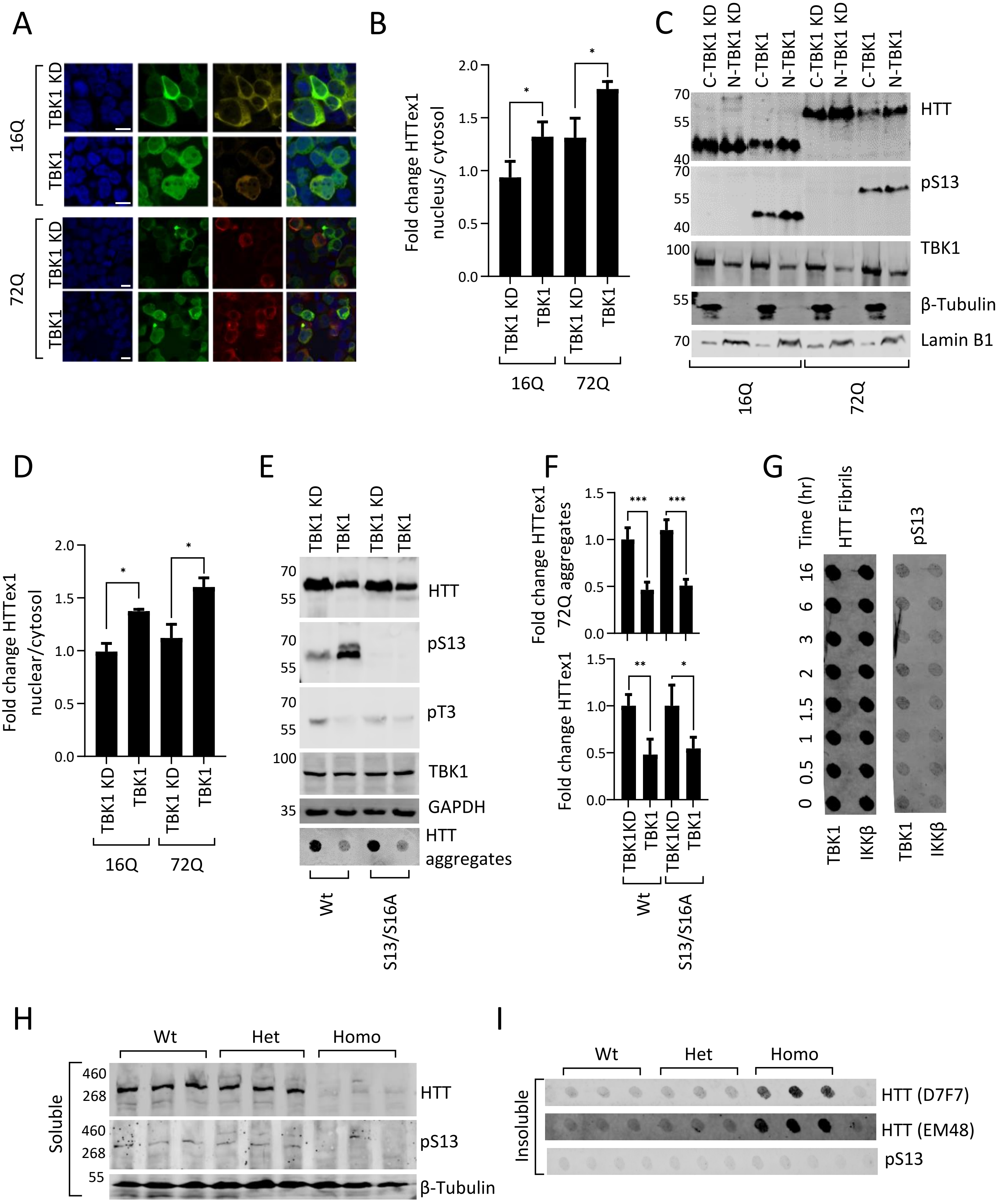
A
16Q
TBK1 KD
TBK1
72Q
TBK1 KD
TBK1
B
Fold change HTTex1 nucleus/ cytosol
TBK1 KD
TBK1
TBK1 KD
TBK1
16Q
72Q
C
C-TBK1 KD
N-TBK1 KD
C-TBK1
N-TBK1
C-TBK1 KD
N-TBK1 KD
C-TBK1
N-TBK1
HTT
pS13
TBK1
β-Tubulin
Lamin B1
16Q
72Q
D
Fold change HTTex1 nuclear/cytosol
TBK1 KD
TBK1
TBK1 KD
TBK1
16Q
72Q
E
TBK1 KD
TBK1
TBK1 KD
TBK1
HTT
pS13
pT3
TBK1
GAPDH
HTT aggregates
Wt
S13/S16A
F
Fold change HTTex1 72Q aggregates
Fold change HTTex1
TBK1KD
TBK1
TBK1KD
TBK1
Wt
S13/S16A
G
Time (hr)
HTT Fibrils
pS13
TBK1
IKKβ
TBK1
IKKβ
H
Wt
Het
Homo
Soluble
HTT
pS13
β-Tubulin
I
Wt
Het
Homo
Insoluble
HTT (D7F7)
HTT (EM48)
pS13

### Sup. Fig. 5

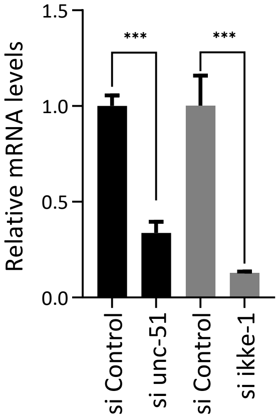
Relative mRNA levels
1.5
1.0
0.5
0.0
***
***
si Control
si unc-51
si Control
si ikke-1

Sup. Fig. 6

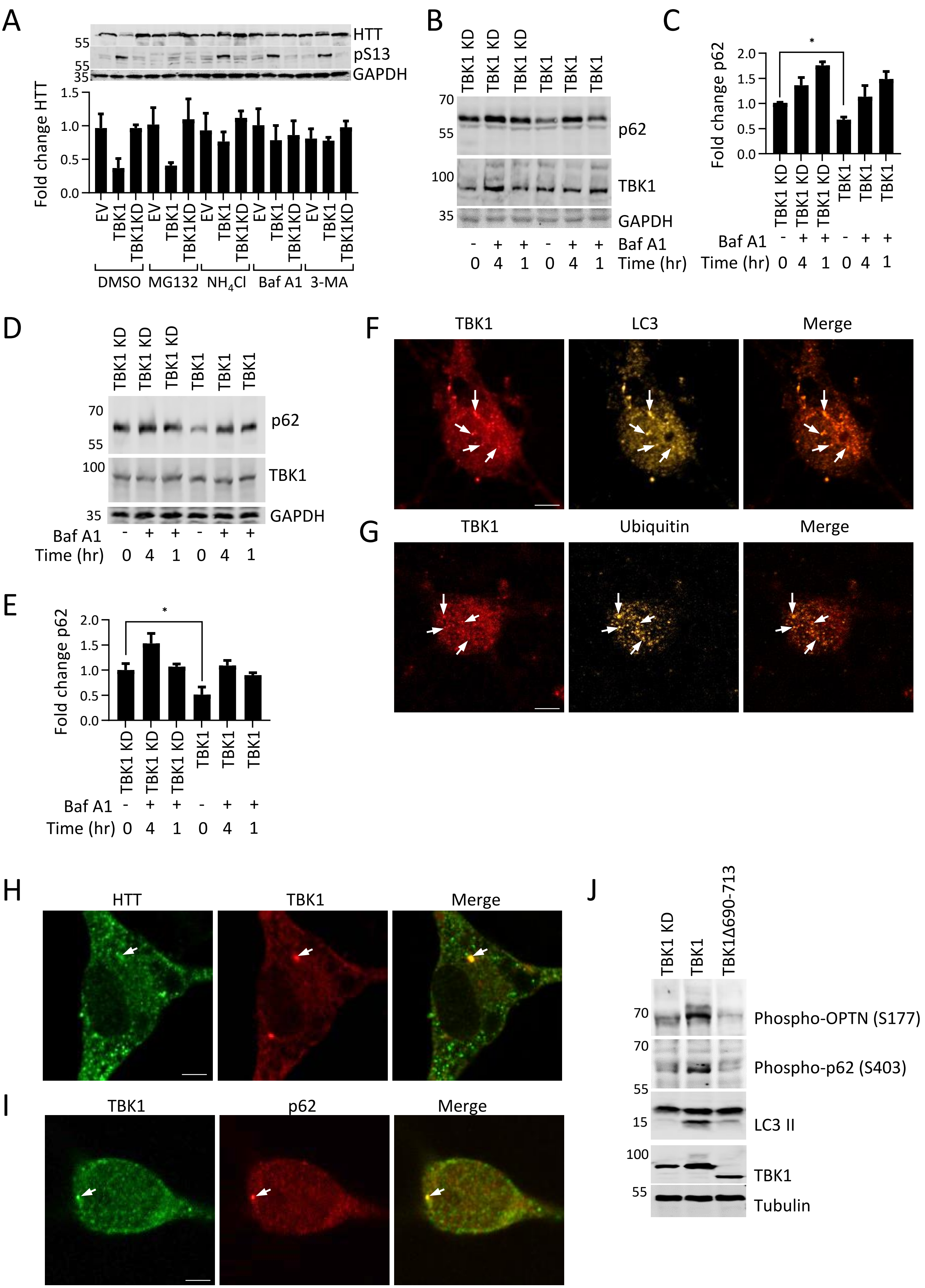
A
HTT
pS13
GAPDH
55
55
35
Fold change HTT
1.5
1.0
0.5
0.0
EV
TBK1
TBK1KD
DMSO
MG132
NH4Cl
Baf A1
3-MA
B
TBK1 KD
TBK1
70
55
100
35
p62
TBK1
GAPDH
Baf A1
Time (hr)
0 4 1 0 4 1
C
*
Fold change p62
2.0
1.5
1.0
0.5
0.0
D
E
F
TBK1
LC3
Merge
G
TBK1
Ubiquitin
Merge
H
HTT
TBK1
Merge
I
TBK1
p62
Merge
J
TBK1 KD
TBK1
TBK1Δ690-713
Phospho-OPTN (S177)
Phospho-p62 (S403)
LC3 II
TBK1
Tubulin
70
70
55
15
100
55

Sup. Fig. 7

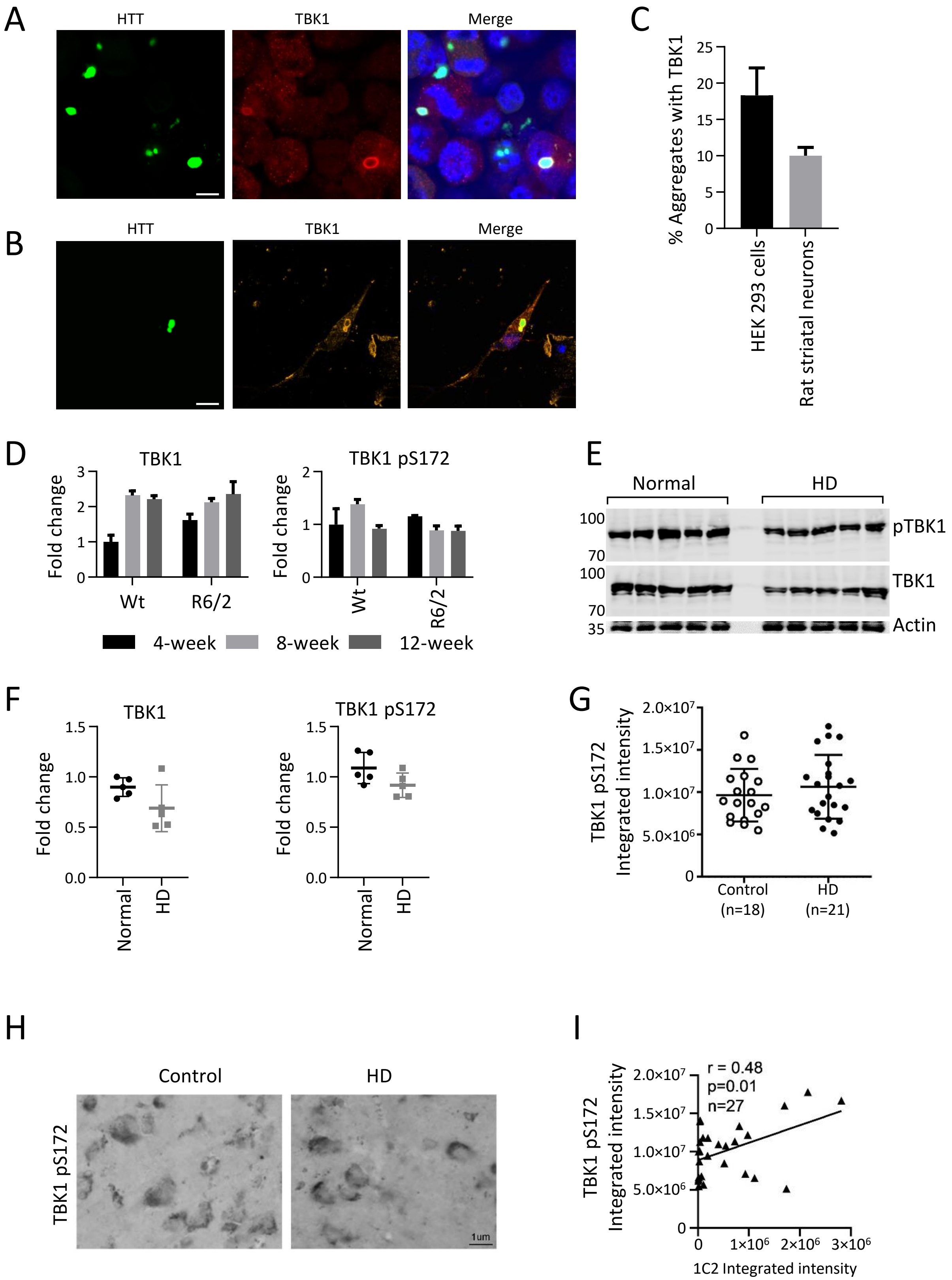
A
HTT
TBK1
Merge
B
HTT
TBK1
Merge
C
% Aggregates with TBK1
25
20
15
10
5
0
HEK 293 cells
Rat striatal neurons
D
TBK1
Fold change
Wt
R6/2
TBK1 pS172
4-week
8-week
12-week
E
Normal
HD
100
70
100
70
35
pTBK1
TBK1
Actin
F
TBK1
Fold change
1.5
1.0
0.5
0.0
Normal
HD
TBK1 pS172
G
TBK1 pS172
Integrated intensity
2.0x10⁷
1.5x10⁷
1.0x10⁷
5.0x10⁶
0
Control (n=18)
HD (n=21)
H
Control
HD
TBK1 pS172
1um
I
r = 0.48
p=0.01
n=27
TBK1 pS172
Integrated intensity
1x10⁶
2x10⁶
3x10⁶
1C2 Integrated intensity
